# Structural insights into 3Fe-4S ferredoxins diversity in *M.tuberculosis* highlighted by a first redox complex with P450

**DOI:** 10.1101/2022.11.02.514812

**Authors:** Andrei Gilep, Tatsiana Varaksa, Sergey Bukhdruker, Anton Kavaleuski, Yury Ryzhykau, Sviatlana Smolskaya, Tatsiana Sushko, Kouhei Tsumoto, Irina Grabovec, Ivan Kapranov, Ivan Okhrimenko, Egor Marin, Mikhail Shevtsov, Alexey Mishin, Kirill Kovalev, Alexander Kuklin, Valentin Gordeliy, Leonid Kaluzhskiy, Oksana Gnedenko, Evgeniy Yablokov, Alexis Ivanov, Valentin Borshchevskiy, Natallia Strushkevich

## Abstract

Ferredoxins are small iron-sulfur proteins and key players in essential metabolic pathways. Among all types, 3Fe-4S ferredoxins are less studied mostly due to anaerobic requirements. Their complexes with <u>cy</u>tochrome <u>P</u>450 redox partners have not been structurally characterized. In the present work, we solved the structures of both 3Fe-4S ferredoxins from *M. tuberculosis* - Fdx alone and the fusion FdxE–CYP143. Our SPR analysis demonstrated a high affinity binding of FdxE to CYP143. According to SAXS data, the same complex is present in solution. The structure reveals extended multipoint interactions and the shape/charge complementarity of redox partners. Furthermore, FdxE binding induced conformational changes in CYP143 as evident from the solved CYP143 structure alone. The comparison of FdxE–CYP143 and modeled Fdx–CYP51 complexes further revealed the specificity of ferredoxins. Our results illuminate the diversity of electron transfer complexes for the production of different secondary metabolites.

## Introduction

Fe-S proteins such as ferredoxins are ubiquitous and ancient proteins indispensable for life^1^. Several lines of evidence suggest that ferredoxins are amongst the oldest proteins on Earth^2,3^. Ferredoxins are responsible for CO_2_ reduction, respiration and other biological electron transfer reactions^4^.

The diversity of the ferredoxin-containing electron cascades is huge. A wide range of reduction potentials can be achieved with Fe-S clusters of various stoichiometries: [2Fe-2S], [3Fe-4S], [4Fe-4S], [3Fe 4S][4Fe 4S], 2[4Fe 4S], each have their own characteristic Fe-S ligating sequence motif and protein scaffold. Reduction potentials of ferredoxins as well as interaction with their cognate redox partners could be further tuned by modifying their amino acid sequences^5,6^, evolving the protein-controlled, energy-conserving electron transfer pathways. Polyferredoxins composed of three to seven 2[4Fe-4S] modules have also been reported^7^.

The [4Fe-4S] clusters are considered as the first to have evolved^8,1^ and ferredoxins of this type are the most ubiquitous and abundantly present in anaerobic organisms, whereas [2Fe-2S] cluster type ferredoxins are abundant in aerobic organisms^9^. The [3Fe 4S] cluster can be considered as a cubane [4Fe 4S] cluster missing one of the irons^10^. This class is found exclusively in bacteria. The [3Fe 4S] clusters can emerge from oxidative damage of [4Fe 4S] clusters; it has been hypothesized as an adaptation to the increased oxygen concentration^11^. Some [4Fe-4S] clusters that contain^11^ a Cys-X-X-Asp-X-X-Cys motif can undergo reversible cluster interconversion to [3Fe-4S]^12^, while most ferredoxins containing [3Fe-4S] clusters do not demonstrate such ability. The factors that control assembly and conversion of these clusters are unknown.

More complex 2[4Fe-4S] clusters are proposed to emerge from the gene duplication^8^.

The explosive data from genome sequencing and metagenome analysis have revealed that bacterial genomes often have a larger number of genes that encode multiple ferredoxins. However, the experimental data on their function and redox partners are largely missing. For example, reduced genome of *M*.*tuberculosis* (Mtb) *H37Rv* encodes five ferredoxins, Fdx (*Rv0763c*), FdxA (*Rv2007c*), FdxC (*Rv1177*), FdxD (*Rv3503c*), and FdxE (*Rv1786*). In addition, it also contains two ferredoxins within fusions, FprB (*Rv0886*) and FdxB (*Rv3554*). In FprB 4Fe-4S ferredoxin domain is arranged at the N-terminus upstream of the reductase domain, while FdxB additionally has a fatty acid desaturase domain at the N-terminus, followed by the ferredoxin reductase domain and a C-terminal 2Fe-2S ferredoxin domain. The function and/or redox partners of these fusions are currently unknown. Such diversity might serve many different physiological functions yet to be defined.

Interestingly, two Mtb 3Fe-4S ferredoxins, Fdx and FdxE, are co-located in respective operons with <u>cy</u>tochrome <u>P</u>450 (CYP) enzymes suggesting their functional relevance. CYPs catalyze a production of primary and secondary metabolites and are involved in the oxidation of a vast range of environmental toxins and drugs. For catalysis, CYPs required electrons typically supplied by redox proteins, such as ferredoxins. Growing evidence has demonstrated that redox proteins can affect the catalytic rate, product profile, the type and selectivity of P450-catalyzed reactions under varied environmental and cellular conditions^13^. Recent comparative analysis of secondary metabolites biosynthetic gene clusters, ferredoxins and CYPs in Bacteroidetes and Firmicutes species from human gastrointestinal microbiota indicate that these two bacterial groups produce different secondary metabolites^14^, which might correlate with their contrasting effects on the human health. However, such systematic analysis is not available for the species belonging to actinomycetes or to the genus Mycobacterium in particular. It is known that nonsterol-producing Mtb Fdx is capable to support CYP51B1 (*Rv0764c*) sterol demethylase activity functioning within the redox chain FprA − Fdx − CYP51B1^15,16^. The ferredoxin *Rv1786* gene is adjacent to CYP143 (*Rv1785c*) whose function is not defined. The binding affinity measured for the pair FdxE–CYP143A1 suggests that they form a redox complex^17^. However, structural details of redox complexes formed by bacterial ferredoxins remain largely unexplored hampering understanding of directional electron transfer from/to cognate ferredoxin partners.

Here we explored genome context for two ferredoxins Fdx and FdxE, determined binding kinetics between FdxE and CYP143, and solved three crystal structures: Fdx, CYP143 and a complex FdxE–CYP143. The complex Fdx –CYP143 represents the first of its kind and we additionally complemented our structural studies by SAXS data in solution. Overall, our experimental work along with AlphaFold2 predictions of an additional relevant redox complex shed light on the specificity of the [3Fe-4S] cluster-containing ferredoxins. Exploring the variety and selectivity of electron carriers in biological processes is crucial for their potential applications as biosensors, biofuel cells, pharmaceuticals as well as in catalysis.

## Results

### Crystal structure of Fdx (Rv0763c)

The ferredoxin Fdx was the first among ferredoxins discovered in Mtb and since then served as an auxiliary redox partner for all known by that time CYPs. CYP51 crystal structure was solved soon after the Mtb genome sequencing^18^. An unusual for mycobacteria steroid demethylase activity of CYP51 was reconstituted with Fdx^15,16^ confirming their redox partnership. Recent progress highlighted a set of different types of ferredoxins in Mtb, but linking them to their cognate redox partners is far from completion. To get structural insight into a single 3Fe–4S type of ferredoxin we determined a crystal structure of Fdx at 2.0 Å resolution (see Table S2). The observed overall fold is typical for the monocluster ferredoxins with two double-stranded antiparallel β-sheets and two α-helices. A short α-helix 1 (Met17 – Glu21), a longer α-helix 2 (Glu43 – Ala55) and two antiparallel β-sheets, a longer β-sheet A (Tyr3 – Ala7 and Leu61 – Glu65) and a short β-sheet B (Phe26 − Agr27 and Glu35 −Ile36) are well defined (Fig. 1A). The structure contains four reverse turns: A, B, C, and E. Turn D is not evident in Fdx forming a loop. The turn C (residues 29 to 33) forms a rather noticeable protrusion from the surface of the protein. The protruding turn is adjacent to the iron-sulfur cluster and considered to be important in specific interactions with electron transfer partners^19^ (discussed below).

**Fig. 1.**
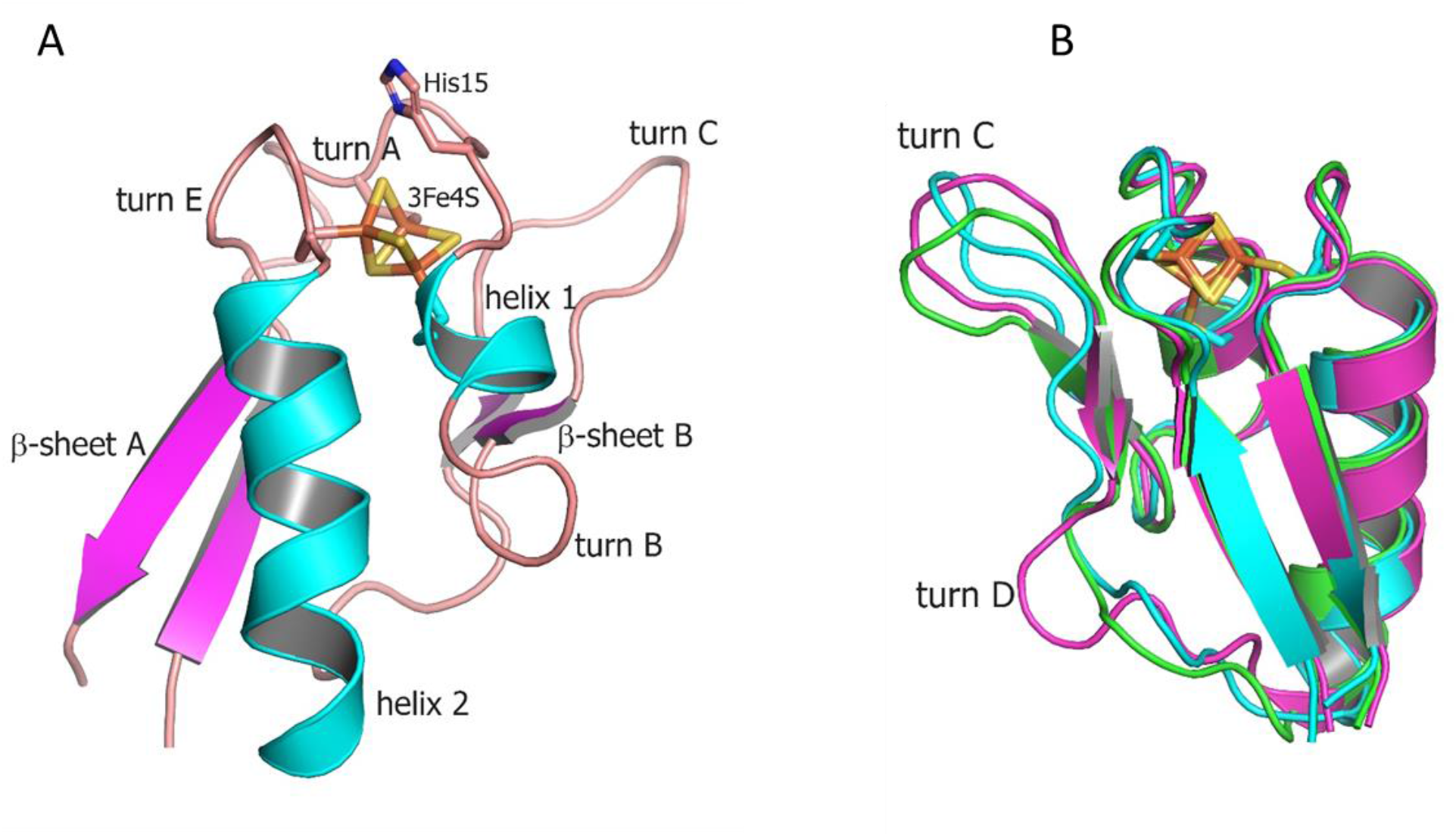
**A-** Overall structure of Fdx ferredoxin of *M*.*tuberculosis*. **B** - Structural alignment of the Fdx (cyan) with other structurally characterized 3Fe–4S ferredoxins from *R. palustris* HaA2 (PDB: 4OV1; magenta) and *P. furiosus* (PDB: 1SJ1; green). The Fe–S clusters are shown in stick representation (Fe, orange; S, gold).

The [3Fe–4S] cluster is bound by cysteine residues 12, 18 and 56. The protein contains no cysteines other than those directly involved in cluster binding. A single [3Fe-4S] cluster geometry is shown in Fig. S1 including values of individual bond lengths and angles for both Mtb ferredoxins (FdxE discussed below). None of the average values for corresponding bonds differs significantly between two solved structures and when compared with other 3Fe-4S ferredoxins. The values for S—Fe—S angles, however, differ between ferredoxins.

Both Fdx and FdxE have the same CXXHXXC(X)nCP motif, where proline is invariantly conserved across all 3Fe-4S ferredoxins (Fig. 2). The His15 residue does not form a hydrogen bond to S2 of the [3Fe–4S] cluster (distance of 3.6 Å); its side chain is turned away (Fig. 1A).

**Fig. 2.**
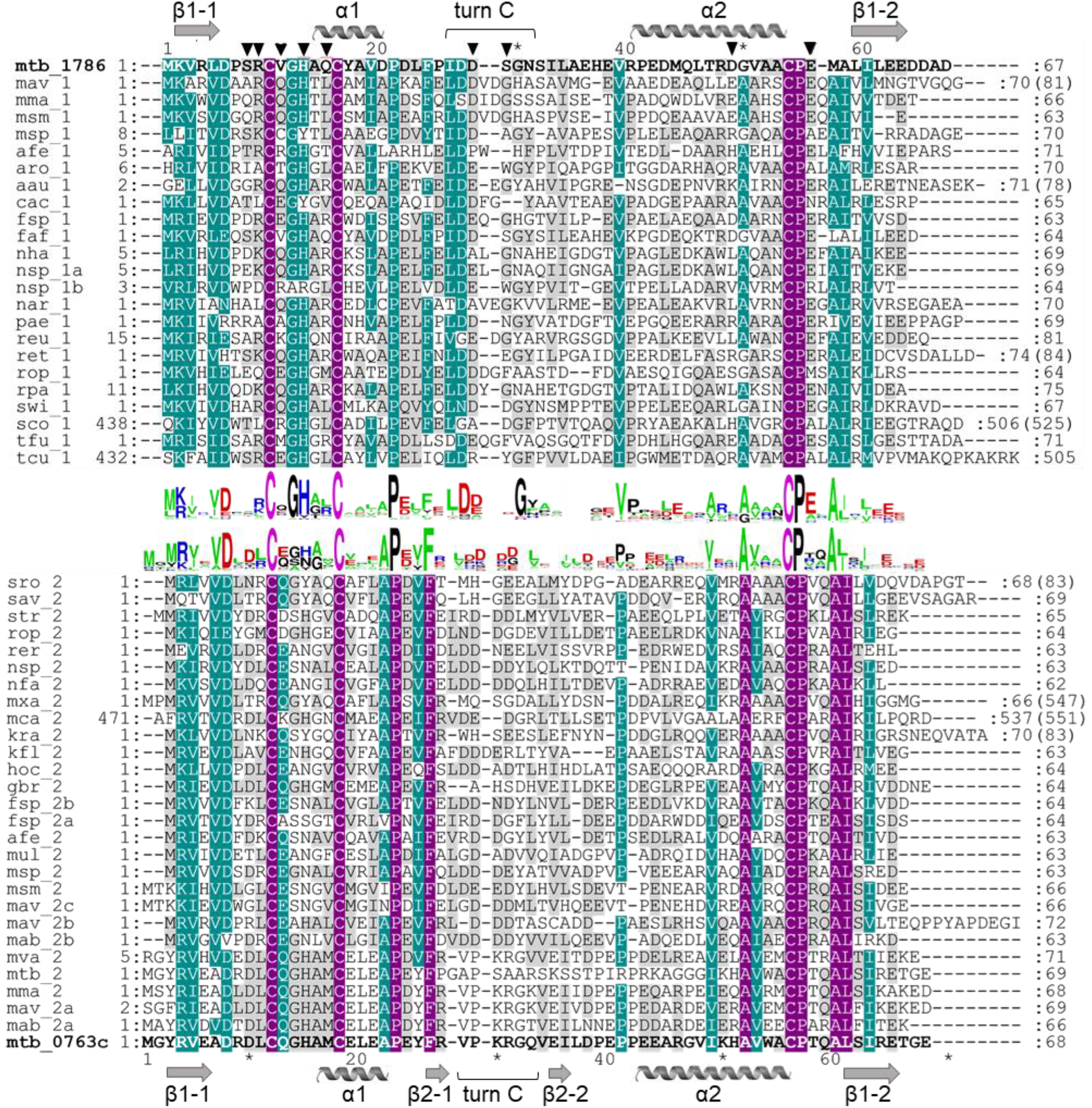
Multiple sequence alignment of ferredoxins taken form the genome context of *Rv1786* (mtb_1786, NP_216302.1) and *Rv0763c* (mtb_0763c, WP_003403890.1) of Mtb (Fig. S3). Description for abbreviation of proteins provided in Fig. S2. Alignment was prepared using Clustal W algorithm. Sequence logos were prepared using WebLogo (https://weblogo.berkeley.edu/). Violet color indicates identical residues, turquoise color - residues with 75% of homology, and grey color - residues with 35% of homology. Numbers at the beginning and at the end of each row correspond to the first and the last amino acids of each sequence taken for alignment. Numbers in brackets indicates the total number of amino acids in the corresponding proteins. Amino acids important for the interaction with redox partner shown as triangles.

From the sequence alignment, it appears that turn C AA composition is specific for Fdx and a small group of related ferredoxins (Fig. 2). Indeed, when the Fdx structure is superimposed to other ferredoxins turn C differs markedly (Fig. 1B). Overall, ferredoxins deduced from the genome context of both Fdx and FdxE (Fig. S3) can be divided to two groups: one is more widely distributed and has classical Asx type (where additionally present a hydrogen bond between a carbonyl oxygen atom of the side-chain of residue n and the amide group of residue n+3) of turn C, while the second group is similar to the Fdx of Mtb and less represented. Furthermore, in the FdxE group species without His residue in the motif do not have CYPs genes in the immediate vicinity. However, in case of Fdx homologs, His might be substituted with N/Y/S (*M. avium, 104, M. abscessus, Haliangium ochraceum, Ktenobacter racemifer* etc), but in gene surrounding it is always CYP. Based on this observation we could suggest that His is not indicative of a redox partner. Further studies will be required to make any correlations.

Homologs of *Rv1786* are present in other species, e.g. in *Frankia sp, Thermobifida fusca* (both have 43% of homology), *Streptomyces griseus* (40%), *Rhodococcus, Nocardioides* (39%), *Rhizobium etli* (34%), *Rhodopseudomonas palustris* (36%), *Arthrobacter aurescens* (33%), *Conexibacter woesei* (34%), *Thermosipho africanus* (33%), *Sphingomonas sp*. (32%)^20^. In most cases, *Rv1786* gene is organized in one operon with respective CYP (Fig. S3A), which might be an evolutionary advantage; however, its function is not yet defined. Exceptions, where *Rv1786* homologs are present alone, could also be found (e.g. *Mycobacterium avium, Rhodococcus opacus)*, but from their different gene environment it is hard to deduce common function. In Mtb CYP143 and *Rv1786* genes are organized within the ESX-5 type VII secretion system implicated in the virulence^21^. Despite recent advances in structural characterization of the ESX-5, the specific substrates translocated by this system are still not identified.

### Interactions between FdxE and CYP143

We used a SPR analysis for direct monitoring of the interaction between FdxE and CYP143. Biotinylated CYP143 was specifically immobilized on a streptavidin-coated chip, while FdxE was used as an analyte. This immobilization strategy ensures that all molecules are in the same orientation and a high protein density is achieved not affecting the potentially interacting sites. Analysis of the kinetic parameters of the complex formation between CYP143 and FdxE (Table 1 and Fig. S4) demonstrated a high affinity binding with K_d_ value in the nanomolar range (55 nM). The FdxE−CYP143 complex is characterized by high on rate (k_on_ ∼ 10^6^ M^-1^s^-1^) and high off rate constants indicating a specific yet transient interaction. The results are in accordance with the biological role of ferredoxins as an electron carrier that shuttle electrons between CYP and reductase. Notably, obtained kinetic constants characterizing the complex formation significantly differ from those previously published^17^. In our experiment, the FdxE−CYP143 complex was formed (k_on_) and dissociated (k_off_) more than 600 and 100 times faster, respectively, than reported^17^, which could be explained by different immobilization strategies and experimental conditions. The main contribution to the high affinity of CYP143 for FdxE is the high rate of complex formation. Such rate of complex formation is comparable with that for antibodies, indicating that the interaction is in diffusion-limited mode^22^. To reduce the contribution of the diffusion effect, we used a relatively high flow rate in our experiments.

**Table 1.**
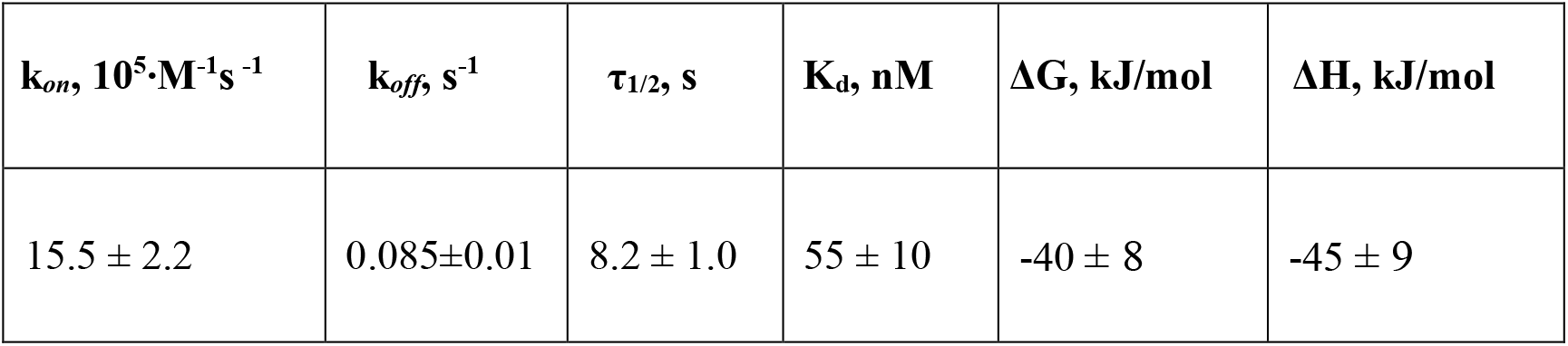
Kinetic and thermodynamic parameters of the complex formation between FdxE and CYP143.

Analysis of the obtained thermodynamic parameters (Table 1) shows that the FdxE−CYP143 complex is enthalpy driven (ΔH<0 value) suggesting the main contributions to complex stabilization are electrostatic interactions and hydrogen bonds^23^. The positive value of the entropy component (-TΔS>0) may indicate the desolvation of polar and charged amino acids upon complex formation and/or the solvent release from the inter-protein region^24^.

### Overall structure of the FdxE−CYP143 complex

Taking into account experimentally determined both high off- and on-rates as well as low stability of the complex (τ_1/2_ = 8.2 s) it seems challenging to trap this transient interaction for structural studies. Here we applied a fusion strategy proven useful in our previous structural studies of the complex between human adrenodoxin and CYP11A1^25^. We constructed a single protein containing FdxE fused via a linker to the N-term of CYP143. This protein was used to obtain a crystal structure of the FdxE−CYP143 complex at 1.6 Å resolution as well as for SAXS studies in solution. In the structure, FdxE binds to the proximal surface of CYP143 and has a typical fold of monocluster ferredoxins as described above for Fdx. CYP143 fold is characteristic for cytochrome P450 and without the ligand presented in the open conformation. The water molecule is coordinating heme iron (Fig. 3). The protein-protein interface is well defined.

**Fig. 3.**
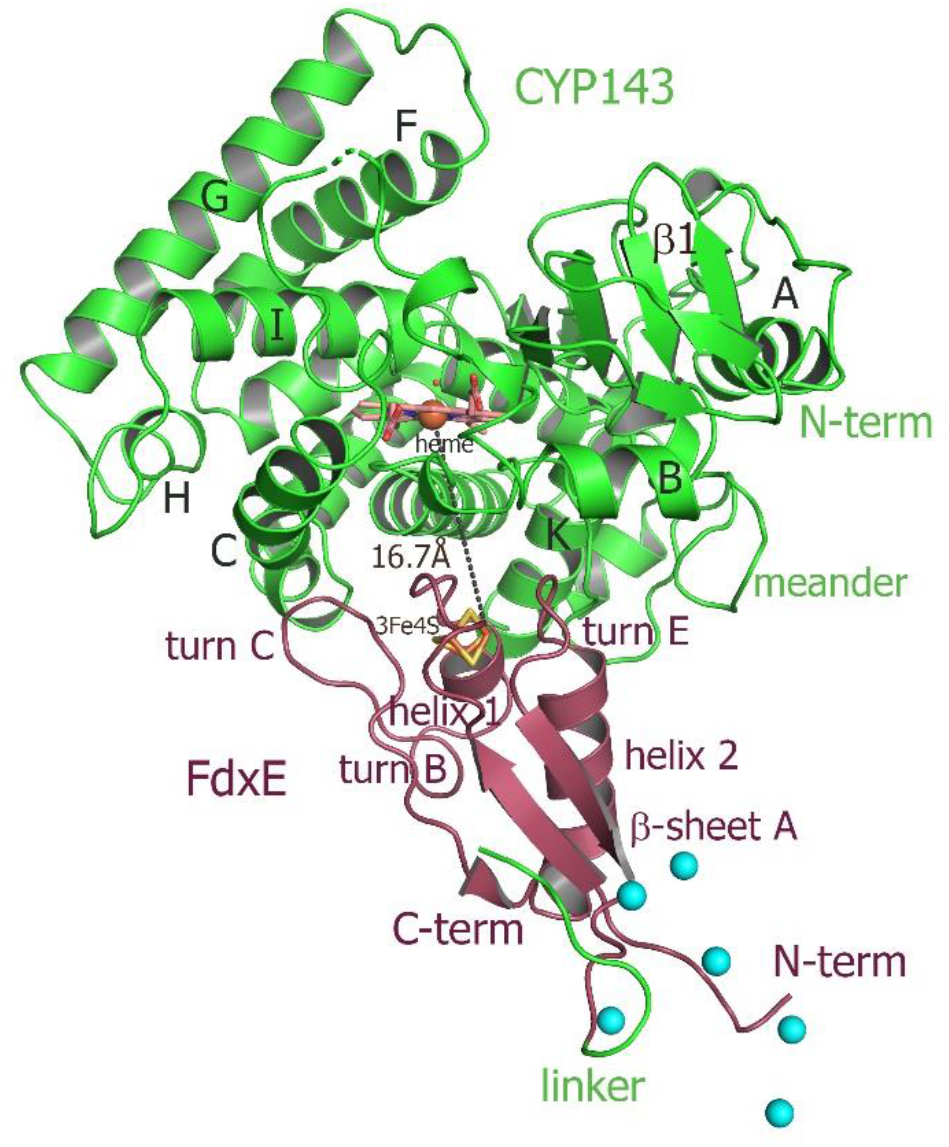
Overall structure of the FdxE–CYP143 fusion protein. FdxE binds (raspberry) at the proximal face of CYP143 (green). Cyan spheres are Ni ions from crystallization conditions bound to His-tag. The N-term His-tag of the FdxE introduced to facilitate purification using metal affinity chromatography is visible and chelated by the Ni ions from crystallization conditions, while the electron density for the part of the linker between two proteins and the first eight residues of CYP143 cannot be resolved.

The buried area upon the complex formation between FdxE and CYP is 1809 Å^2^. The shortest distance between the CYP143 heme group and the ferredoxin iron–sulfur cluster is 16.7 Å. As expected, several structural regions from the CYP143 proximal side are participating in a complex formation, namely helices C, K and L and the meander preceding the heme-containing loop. However, some specifics of the 3Fe–4S ferredoxin−CYP complex can be identified. First, the angle of FdxE approaching CYP, second, a more extended interaction interface (additionally involving B helix of CYP and turn С of FdxE), and third, the shape/steric complementarity. Of the latter, particularly interesting is the complementarity between the helix 2 of FdxE and CYP meander running along each other; between the helix 1 of FdxE and K helix of CYP; a positioning of the heme coordinating loop right above the [3Fe–4S] cluster and finally, a perfect fit of the turn C of FdxE between С and D helices of CYP. Altogether, it indicates a high specificity of this type of electron transfer complex.

### Interactions between FdxE and CYP143

The electron density of the iron–sulfur cluster of FdxE clearly identifies a [3Fe–4S] cluster coordinated by Fe—S bonds to cysteines Cys10, Cys16 and Cys54. The [Fe3–S4] cluster has typical cuboidal geometry with similar ∼2.3 Å bond lengths between the cysteinyl sulfur and Fe (Fig. S1). A residue His13 of the CXX**H**XXC(X)nCP motif is not ligated to the [3Fe–4S] cluster (Fig. 4). Instead, the N1 atom of the imidazole ring of His13 forms a hydrogen bond with the O^Σ1^ atom of Glu56 from the opposite loop around cluster. The other interactions of His13 are water mediated contacts with the CYP143 residues: the N3 atom of the imidazole ring - with Arg349 of L helix, and the main chain oxygen - with Arg266 from K helix.

**Fig. 4.**
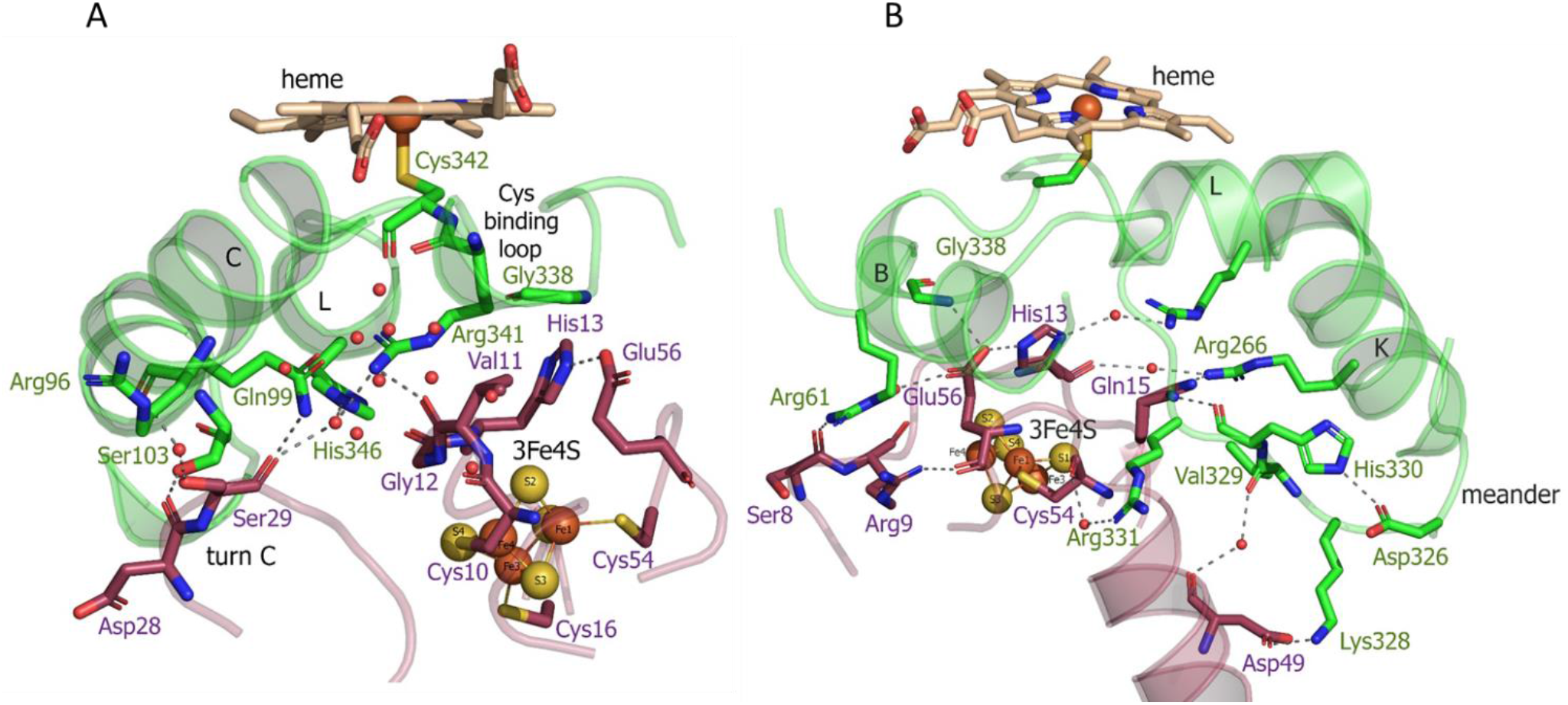
Two views of the interactions between FdxE and CYP143.

The side chain of Glu56 of FdxE is in direct contact with Gly338 of the Cys-coordinating loop of CYP and involved in a water-mediated interaction with the side chain of Arg61 from the B helix of CYP143. The main chain of Glu56 is stabilized by the internal interaction with Arg9 that precede cluster-coordinating cysteine. The CYP residue Arg61 also forms a weak hydrogen bond (distance of 3.1 Å) with the main chain oxygen of Ser8 in turn A of FdxE. In a complex, His13 and Glu56 residues are positioned right below the Cys-coordinating loop of CYP.

In addition, the complex is stabilized by the salt bridge between Arg266 of CYP143 K helix and Gln15 preceding the cluster coordinating Cys16 of FdxE (Fig. 4B). Although the side chain of Gln15 modeled in two conformations, it does interact with CYP, either with Arg266 or with the main chain of His330 from the meander.

A meander region of CYP is sandwiched between a K helix of CYP and Helix 2 of FdxE. The inner salt bridge in the meander, between residues Asp326 and His330, maintains its specific conformation so that Lys328 forms a salt bridge with Asp49 of FdxE (Fig. 4B). Additionally, this interaction region is stabilized by two water-mediated contacts: between the main chain oxygen of Asp49 of FdxE and the main chain oxygen of Val329 of CYP, and between the main chain oxygen of Cys54 coordinating cluster of FdxE and side chain of Arg331 of CYP.

The loop coordinating heme of CYP is adjacent to the meander region and involved in the interaction with the redox partner. Here, in addition to above mentioned Gly338, the residue Arg341 immediately preceding the proximal ligand of heme is H-bonded to Val11 of FdxE.

Finally, the interaction spot between redox partners is also observed on the periphery of the interaction interface (Fig. 4A). The turn C of FdxE perfectly fits between two adjacent helices C and D of CYP, forming a hydrogen bond between Asp28 of FdxE and Ser103 of CYP. The intermolecular interaction is additionally stabilized by a water-mediated interaction of Ser29 of FdxE and Arg96 of CYP. The same Ser29 also forms a weak H-bond (distance is 3.1Å) with Gln99 of CYP and water mediated contacts with Arg341 and His346 of CYP. Notably, the water chain observed in this region is filling the interaction interface between the loop harboring the Cys10 ligand of the [3Fe-4S] cluster and the C helix of CYP.

Overall, the hydrogen bonding interactions predominate in the formation of the FdxE–CYP143 complex consistent with the obtained thermodynamic data.

Based on the distance to the heme, two irons of the [3Fe-4S] cluster, Fe4 (16.7 Å) and Fe1 (16.8 Å) might participate in the electron transfer. Electrons could flow from the Fe4 of the cluster to the Cys10 –Val11 peptide or, alternatively, from Fe1-S2 to the Gly12 – Val11 peptide and then via Arg341 and cysteinyl ligand (Cys342) directly to the heme iron. The through-space jumps via side-chain atoms of His13 and Arg341 cannot be excluded. The precise positioning of the cluster right below the heme-binding loop, the short, efficient and coupled pathway for electron flow to the heme iron, indicates that the complex between two proteins is specific.

### CYP143 structural changes for complex formation

To visualize structural changes in CYP143 induced by the interaction with FdxE we solved the CYP143 structure alone at similar resolution (1.4 Å). It is worth mentioning that attempts to crystallize FdxE alone were unsuccessful. CYP143 structure is conforming to the typical cytochrome P450 fold with 12 α-helices (A - L) and 4 β-sheets. In the absence of the ligand protein crystallized in open conformation with the heme cofactor accessible to the bulk solvent. The iron of the heme is hexacoordinated by the invariant Cys342 and a water molecule. This water is stabilized by the H-bond with glycerol molecule from a cryoprotectant solution. Superposition of the CYP143 structure with the CYP143 part of the complex structure shows RMSD = 0.932Å for all atoms and reveals a different position of the lower part of the meander (Fig. 5A). In the CYP143 structure alone, the meander is away from the K helix, while in the complex it moves closer and sandwiched between helix 2 of FdxE and K helix of CYP. The relocation distance of 5.5 Å is observed for the Gly327 Ca atom. Upon complex formation the salt bridge is formed between Asp326 and His330 shaping the meander for the interaction with ferredoxin by a salt bridge between Lys238 of CYP and Asp49 of FdxE (Fig. 5A) as well as water-mediated contacts described above.

**Fig. 5.**
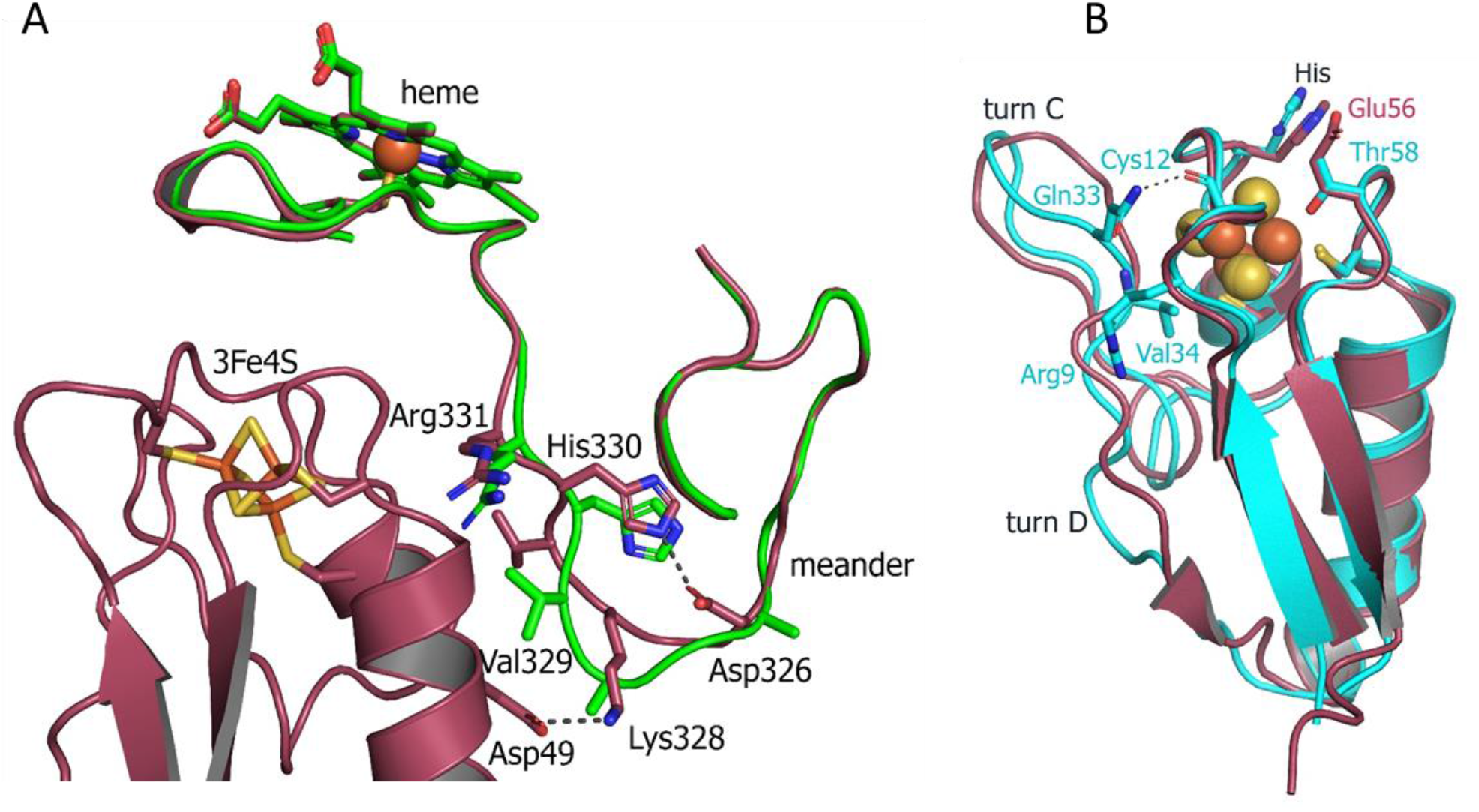
**A-** Superposition of CYP143 structure alone (green ribbon) with the structure of the FdxE−CYP143 complex (raspberry) detailing the meander region. **B** - Superposition of Fdx (cyan) and FdxE (raspberry).

A closer examination of interacting regions of CYP143 reveals different side chain conformation of the charged residues involved in intermolecular contacts with Fdx in a complex structure − Gln99 of C helix, and Arg266 of K helix.

With the CYP143 structure alone in hands, we calculated its electrostatic surface potential (Fig. S5A). The local proximal area involved in the interaction with FdxE is positively charged, despite overall theoretical pI=6.8. With FdxE being negatively charged (Fig. S5B) one could suggest electrostatic steering of redox partners when they approach each other.

### Small-Angle X-ray Scattering Analysis (SAXS)

To obtain the information about the structure and conformational flexibility in solution the fusion complex FdxE−CYP143 was investigated by SAXS. Assuming that CYP143 and FdxE positions are fixed at their crystal positions, we performed SAXS-based modeling using CORAL program^26^. Obtained structure (Fig. 6) approximates well SAXS data (χ^2^ = 1.41, Fig. S6A and Table S2). It strongly indicates that the FdxE−CYP143 complex in solution is similar to that in the crystal structure, and, consequently, the contacts between CYP143 and FdxE are not an artifact of the crystallization.

**Fig. 6.**
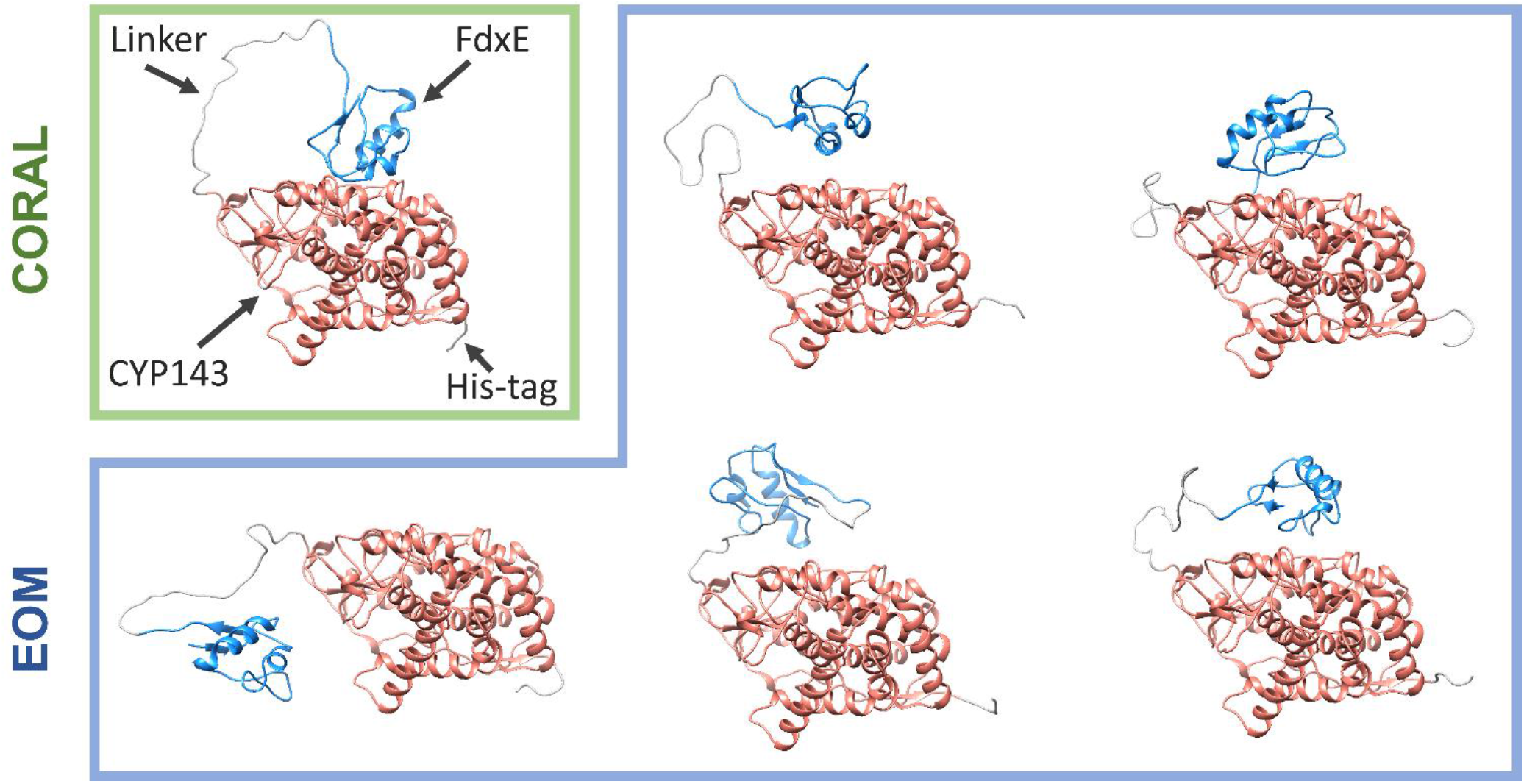
Results of SAXS data analysis of the fusion FdxE−CYP143 complex. The models obtained with CORAL and EOM. The models aligned to the same position of CYP143.

Additionally, we used Ensemble Optimization Method (EOM)^27^ to fit SAXS experimental data with an ensemble of proteins with variable positions of CYP143 and FdxE. EOM is suitable for describing the distribution of the relative position of two domains connected by a linker, even if these domains differ significantly in molecular weight^28^. EOM generated a large pool of possible conformations (that has a distribution of *R*_*g*_shown in Fig. S6B) and then it selected a subset of conformations that minimizes the discrepancy χ^2^ between theoretical and experimental SAXS data. The distribution of *Rg* for the selected subset of conformations corresponds to the smallest sizes compared to the initial pool of conformations, indicating that the majority of the protein molecules in solution are in the most possible compact conformations.

The multistate model generated by EOM (Fig. 6), which delivered the best fit of SAXS data (χ^2^ = 1.16), contains four out of five structures similar to the position of FdxE in the crystal structure. The fifth structure of the multistate model is more expanded and clearly demonstrates that the linker length and flexibility enable free movement of FdxE. The multistate model indicates that a dynamic equilibrium takes place in the solution, and about 80% of the proteins form complexes that are similar to crystal structure.

### Comparison of Fdx and FdxE structures

To understand the specificity of two ferredoxins from Mtb we compared their structures. Although both ferredoxins have the same fold and share the same cluster coordinating motif some differences are clearly seen. The structures were aligned with Cα RMSD = 1.614 Å. Two main differences are - the absence of turn D in Fdx and the position of turn C (Fig. 5B). The latter is of particular importance as it is involved in the interaction with a redox partner. As mentioned above, not only the conformation but the amino acid composition of the turn C differ between two ferredoxins. In the Fdx structure, a turn C is additionally stabilized by the interactions with the turn A residues: backbone Val34 with Arg9 and Gln33 with backbone Cys12. The His13 residue from the motif interacts with Glu56 residue in FdxE structure, while the corresponding Thr58 residue in Fdx is pointed towards the cluster with the distance to the S2 = 3.7 Å.

The residues interacting with the redox partner are also different between two ferredoxins. Specifically, Val11 implicated both in the interaction with the redox partner and electron flow corresponds to Gln13 in Fdx. The surface residues Ser8, Arg9 from A turn and Gln15 from helix 2 of FdxE correspond to Asp10, Leu11 and Met17 in Fdx, respectively (Fig. 2). So the polar contacts in these regions are opposite in charge for two ferredoxins. In addition, Asp49 in Fdx that forms a salt bridge with Lys238 of the adjusted meander of CYP replaced by His51 in Fdx further revealing their specialization.

## Discussion

In this work, we focused on two 3Fe-4S ferredoxins from Mtb associated with CYPs. We solved crystal structures for both of them, one is for ferredoxin alone and a second is in the complex with CYP143 (Table S2). For better understanding of the redox partner induced structural changes we solved a crystal structure of CYP143 alone. With this set of structures, we noticed the rigid structure of small ferredoxins, but adaptations for the interaction seen in CYP.

We explored the genetic context for both ferredoxins. Evolutionary advantages may be gained by colocalizing redox partner genes into the same operon and/or adjacent to each other. A 3Fe 4S ferredoxins of *Mycobacterium marinum*^29^, *Streptomyces griseolus*^30^ and *Rhodopseudomonas palustris*^31,32^ are associated with cytochrome P450 enzymes with demonstrated in vitro catalytic efficiency.

In Mtb only two ferredoxin genes are organized in this manner, albeit with notable difference: the reading frame for *Rv1786* is in the opposite direction relative to *CYP143* (Fig. S3A), while it is the same for *Rv0763c* and *CYP51* (Fig. S3B). Of note, in other species homologs of *Rv1786* are organized in one direction within the same operone with *CYP*, indicating functional relevance (Fig. S3A).

In general, CYP51−Fdx fusions are not rare in nature and can be found in different bacterial phyla^33,34^ including Actinobacteria to which Mtb belongs. Native fusion McCYP51−Fdx from *Methylococcus capsulatus* has 3Fe-4S type of ferredoxin and was hypothesized to emerge due to the mutation of a CYP51 nonsense codon leading to translational read-through to an adjacent ferredoxin^34^. The interactions between redox partners within McCYP51−Fdx were described as transient with various orientations of ferredoxin molecule^35^. Of note, CYP51−Fdx fusions still require a reductase component for catalysis. In case of *Rv0763c* homologs (Fig. S3), the fusion with reductase was observed in *Myxococcus xanthus DK 1622* further expanding the fusion capacity of ferredoxins.

We were unable to find CYP143-related fusions with *Rv1786*. However, the search using CXXHXXC(X)nCP motif revealed a native fusion in mercury methylating marine bacteria^36^ which is distantly related to Rv1786 (30.85% homology), where the N-term ferredoxin domain is fused to CYP. Overall, the fusion strategy is common in bacteria, including pathogenic species. Some relevant examples of fusions are discussed below.

Phthalate Family Oxygenase Reductase-like fusion enzymes (abbreviated as PFOR) comprising of CYP116 family are self-sufficient, e.g. NAD(P)H binds directly to the reductase domain and electrons are transferred through FMN, then the 2Fe–2S center and onto the P450 heme iron. CYP116 PFOR fusion proteins are found in *Rhodococcus, Burkholderia, Ralstonia, Labrenzia, Acinetobacter, Toriphiles* and other species. Recently the 3D-structure of CYP116B46 from *Tepidiphilus thermophilus* was solved^37^ revealing the overall arrangement of the redox chain. In the structure, 2Fe–2S ferredoxin domain is oriented towards the reductase domain with the distance between FMN and 2Fe-2S cofactors 7.9 Å, which represents the initial arrangement for the electron transfer. The ferredoxin domain must have sufficient mobility upon reduction for the next step where it interacts with the CYP domain. The molecular dynamic simulation studies confirm the transit of ferredoxin from “distal” to “proximal” conformation enabling efficient electron transfer from the reduced ferredoxin to heme domain^38^. However, there is still the possibility of a dimeric complex (as for P450BM3^39^) with the inter-monomer electron transfer. Of note, the complex between ferredoxin domain and CYP domain in CYP116B46 was modeled using the reference structure of the designed fusion of native redox partners, CYP11A1 and adrenodoxin, we solved earlier^25^. Up to date despite the ferredoxins biodiversity in CYP-mediated reactions, the crystal structures of complexes with CYPs are limited to 2Fe-2S ferredoxins.

In this work we designed a fusion FdxE−CYP143 of Mtb using linker from the PFOR of *Rhodococcus sp. NCIMB 9784* which we used for the fusion of mammalian redox partners^25^. The linker acts as a flexible hinge allowing interaction between CYP and ferredoxin. Using this strategy, we were able to characterize the FdxE−CYP143 complex and highlight certain features related to the 3Fe-S4 type of ferredoxin. However, how the specificity of ferredoxins in CYP-mediated reactions realized and controlled in different bacteria is still an open question. Is there any correlation between the motif ligating cluster and ferredoxin function/redox partners? Could ferredoxins with the same motif substitute each other in different conditions the organism may encounter? To address simpler latter question we modeled the CYP51–Fdx complex using AlphaFold2^40^ and compared it with our experimentally obtained structure of the FdxE−CYP143 complex (Fig. 7).

**Fig. 7.**
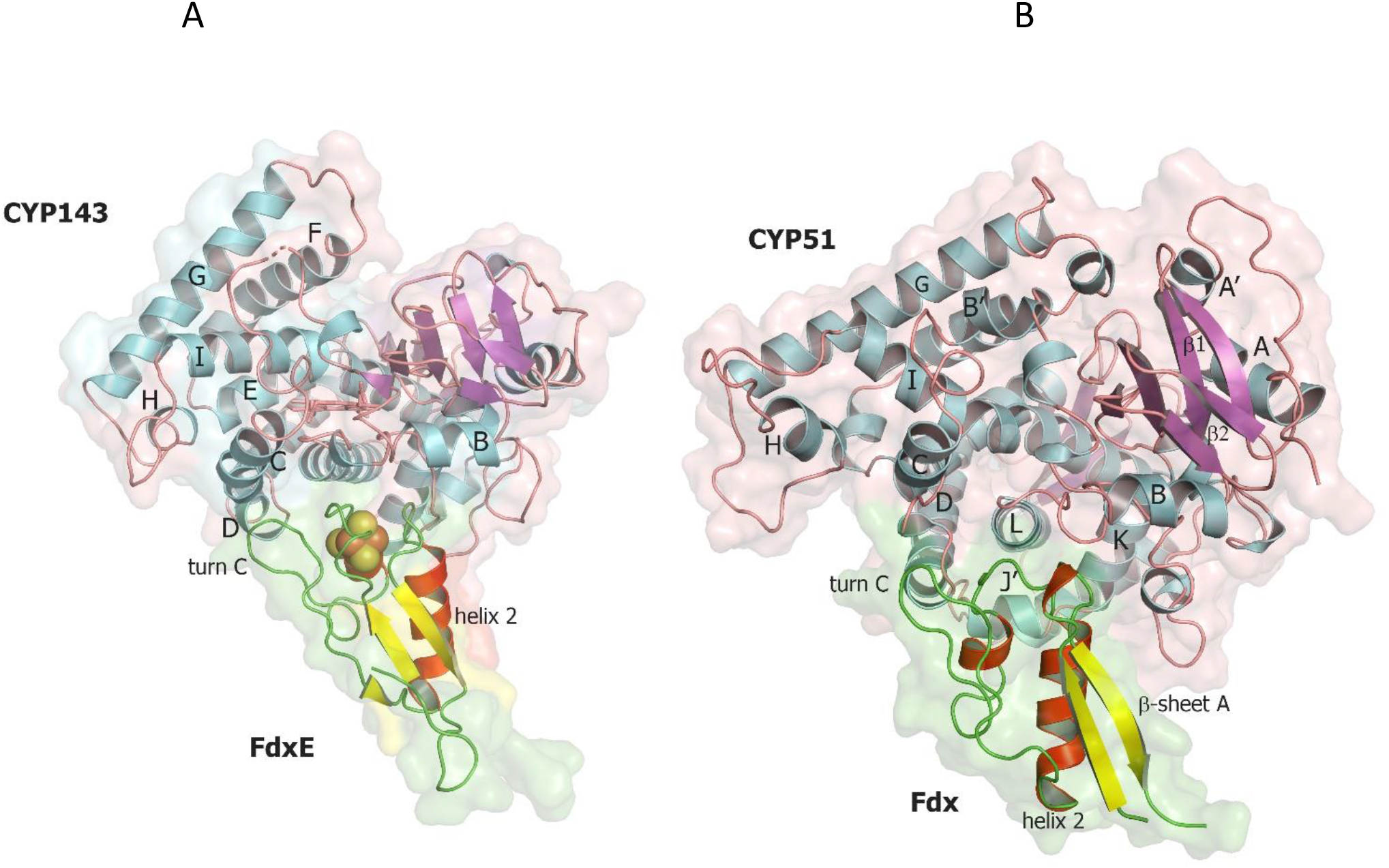
Comparison of two complexes of CYP with 3Fe-4S ferredoxins. A. FdxE−CYP143 complex obtained in this work, B. CYP51̶ Fdx modeled with AlphaFold2.

In the modeled CYP51–Fdx complex the position of Fdx is shifted to the C helix so that Cys-coordinating loop is not right above the cluster, but also shifted and above the turn C of Fdx (Fig. 7B). CYP51 proximal surface is strikingly different from CYP143 as MtbCYP51 structure has quite unique features: a bent I helix and an open conformation of BC loop^18^ and as a consequence the repositioning of the C- and H-helices and the adjacent loops. The observed docking position of the Fdx ferredoxin could also be related to its unusual turn C conformation affecting interaction with CYP. In addition, CYP51 has J′ helix missing in the CYP143 structure. These structural differences along with amino acid sequence differences between Fdx and FdxE (Fig. 2) led to the deviations in the interacting residues between redox partners. One can speculate that conformational changes would take place on the proximal surface of CYP51 upon Fdx binding and CYP51 fold would be more similar to other CYP51 homologs. A recently reported CYP51 structure from *Mycobacterium marinum*^41^ is similar to MtbCYP51 suggesting specificity of bacterial cognate ferredoxins.

In cytochrome c reductase activity assay Fdx and FdxE demonstrated different preferences for the reductases^42^. Furthemore, FdxE is significantly less sufficient in supporting in vitro catalytic activities of Mtb CYP124A1 (^42^ and our own unpublished data), CYP125A1, and CYP142A1 and does not support CYP121 catalysis^42^ suggesting it is less promiscuous and more specific than Fdx. A summary of the *Rv0763c* and *Rv1786* gene expression using MicrobesOnline^20^ also reveals that they are induced and downregulated at different stress conditions (Fig. S8). From this analysis it appears that *Rv0763c* and *Rv1786* cannot substitute each other in supporting respective CYP-mediated reactions, that is feasible considering their organization in the *M. tuberculosis* genome. Further studies should be focused on the search of potential substrates for CYP143 to get more information about the regulation of interaction between these redox partners.

In summary, many ferredoxin genes associated with the CYPome in bacteria provide more specific control over synthesized metabolites under different conditions. However, these systems have yet to be characterized in detail. We believe that our data shed light on the structural preferences driving protein-protein interactions within these electron transfer complexes.

## Methods

### Cloning, expression and purification of Fdx, FdxE, CYP143, and fusion

We amplified the Fdx coding sequence with 6-histidine tag on C-terminus using atcatatgggctatcgagtcgaagc forward primer and taggatccttaatggtgatggtgatggtgctctcccgtttctcggatg reverse primer. Mycobacterium bovis BCG genomic DNA was used as a template instead of M. tuberculosis, since the Fdx sequences are identical in both genomes. The PCR product then was cloned into a pET11a vector using NdeI and BamHI restriction enzymes. The insert sequence was validated by sequencing. The FdxE was cloned similarly into the expression vector pET11a.

The Fdx was expressed in *E. coli* BL21 (DE3). The cells, harboring the expression vector, were grown in TB medium at 37 °C with addition of 100 ug/mL ampicillin and 0.5 mM FeCl_3_. When the culture reached OD_600_ ∼0.7, Fdx expression was induced with 0.7 mM IPTG. The cells were harvested after incubation at 22 °C for 16 hours post induction. The cells were resuspended in buffer A (50 mM TrisHCl pH 7.5, 300 mM NaCl, 20% w/v glycerol) containing 1 mM PMSF and disrupted with Emulsiflex C3 Homogenizer. The lysate after centrifugation was applied on Ni-NTA agarose column, followed by washing step with buffer A containing 60 mM imidazole, and gradient elution with buffer A containing 500 mM imidazole. Fractions with absorption peak at 412 nm were diluted 30-fold with buffer B (10 mM TrisHCl pH 7.5, 20% w/v glycerol) and applied on the DEAE-Sepharose column. After a 20 CV washing step with buffer B, the protein was eluted with linear gradient of buffer C (50 mM TrisHCl, 1 M NaCl, 20% w/v glycerol). Fdx concentration was calculated using molar extinction 11300 M^-1^cm^-1^ at 412 nm ^15^.

The FdxE was expressed in *E. coli* JM109. Overnight culture (3 ml) was used to inoculate 0.5 liter of TB-medium containing 100 mM potassium-phosphate buffer, pH 7.4 and ampicillin (100 µg/ml). The mixture was incubated in a thermostatic orbital shaker at 37°C and 180 rpm. After reaching OD_600_ ∼ 0.4, FdxE expression was induced by adding IPTG (0.5 mM), ampicillin (100 µg/ml). A solution of FeCl_3_ (100 µg/ml) was also added at this point. After 24 h of incubation at 26°C and 100 rpm, the cells were collected by centrifugation (8000g, 10 min). The pellet was resuspended in a 50 mM potassium phosphate buffer, pH 7.4, containing 20% glycerol, 0.1 mM EDTA, 0.5 mM PMSF. Cell suspension was sonicated in an ice-water bath (7×1-min pulses with 1-min intervals). The suspension was centrifuged for 1 h at 20500 rpm and the supernatant was applied to a column with DEAE-sepharose equilibrated with buffer A (50 mM potassium phosphate buffer, pH 7.4, containing 0.1 mM EDTA). The column was washed with 2-3 volumes of buffer A, and then with 10 volumes of buffer A containing 15 mM NaCl. FdxE was eluted from the column with buffer A, containing 200 mM NaCl. Eluted fractions were applied to Superdex 200 16/60 column equilibrated with buffer A, containing 200 mM NaCl. The colored fractions containing FdxE were collected.

The constructs for CYP143, CYP143-Avi-tag at C-terminus and FdxE−CYP143 fusion were cloned in pCW-LIC vector by LIC and co-expressed with GroEL-GroES in *E. coli* strain DH5a in Terrific broth medium at 26 °C with shaking at 100 rpm for 48 hours after induction. In the case of CYP143-Avitag, TB medium was supplemented with biotin (50 µM). Induction was performed at OD_600_ ∼ 0.8 by IPTG (0.5 mM) and arabinose (2 g/L). 0.65 mM δ-aminolevulinic acid was added to the growth medium as heme precursor. For FdxE−CYP143 fusion, FeCl_3_ (20 µM) was added additionally. Cells were collected by centrifugation (3500 rpm for 20 min at 4 °C) and resuspended in 50 mM Tris-HCl pH 7.4 buffer, containing 0.3 M NaCl, 0.5 mM PMSF.

FdxE−CYP143 fusion was purified using metal affinity and ion-exchange chromatography. Cells were lysed by passing through Emulsiflex C5 Homogeniser (Avanti, Canada) twice and then centrifuged to remove membrane fraction (22 000 rpm, 4°C for 1 hour). Supernatant was loaded on an IMAC column (5 ml Ni-NTA His-Trap HP, Cytiva). Column washed with 15 volumes of buffer 50 mM Tris-HCl pH 7.4, 0.3 M NaCl, 25 mM imidazole, and the protein was eluted with a linear gradient of imidazole (0.025 – 0.5 M). Red fractions were pooled and purified on Q-Sepharose column (Source 30Q, Cytiva) pre-equilibrated with buffer 10 mM Tris-HCl, pH 7.4. After washing the column with 15 volumes of buffer containing 0.1 M NaCl, the protein was eluted with a linear gradient of NaCl (0.10 – 1 M). The protein fractions were analyzed by SDS-PAGE and spectrophotometrically, then pooled and concentrated; glycerol was added for storage. CYP143 was purified similarly. Biotinylated CYP143 was purified using affinity chromatography followed by dialysis. Proteins were frozen in liquid nitrogen and stored at -80 °C.

### Crystallization of Fdx, CYP143 and FdxE−CYP143

For Fdx crystallization we used a sitting drop method. Crystals grew at 20 °C in drops containing 1 ul of the protein at concentration 9 mg/ml and 1 ul of 0.1 M Tris-HCl pH 8.5, 0.2 M MgCl_2_, and 25% w/v PEG 3350.

CYP143 (150 µM) and FdxE−CYP143 (200–250 µM) were crystallized in 96-well plate using a sitting-drop method with commercially available kits from Qiagen (NeXtal Classics II screen) and Molecular Dimensions (Structure screens 1 and 2) at 20 °C with 1:1 protein/mother liquor ratio with the ligand concentration of 1-2 mM. Red colored crystals of CYP143 appeared overnight and FdxE−CYP143 in 2 weeks. The best crystals of CYP143 and FdxE−CYP143 grew in 0.2 M Sodium chloride, 0.1 M Bis-Tris pH 6.0, 25% (*w/v*) PEG3350.

### Surface Plasmon Resonance

SPR analyses were carried out using the optical biosensors BiacoreX100, Biacore 3000, Biacore 8K (Cytiva, Marlborough, MA, USA) and sensor chips of SA series S type (Cytiva, Marlborough, MA, USA) at 25 °C. The buffer PBS (10 mM Na_2_HPO_4_, 1.8 mM KH_2_PO_4_, 137 mM NaCl, 2.7 mM KCl, pH 7.4) (Cytiva) was used as a running buffer for CYP143 immobilization and SPR analysis. Before immobilization the chip surface was conditioned with 1M NaCl, 50 mM NaOH solution injection at a flow rate of 30 ul/min for 1 min. Next, 480 nM solution of biotinylated CYP143 with AVI-tag in PBS buffer was injected into the working channel of the biosensor for 7 min at a flow rate of 10 μL/min. The mean final level of immobilization was 5500 ± 500 RU. Reference channel without immobilized CYP143 was used to correct the effects of the non-specific binding of analytes to the chip surface.

### Estimation of kinetic and equilibrium parameters

To assess the parameters of protein–protein interactions, FdxE was injected at various concentrations (5 to 75 nM) through the cell with CYP143 immobilized on the SA chip. The resulting sensorgram represents the difference between the experiment (with immobilized CYP143) and the control (without CYP143) channels. The flow rate was 30 μl/min, the contact time was 2 min. No regeneration of the chip surface was required as the complex dissociates completely within 2 minutes. The resulting sensorgrams were processed in the BIAevaluation 4.1.1 software using the 1:1 interaction model with mass transfer. The τ_1/2_ values characterizing half-time dissociation of a protein-protein complex were calculated from k_off_ values according to the equation (1).

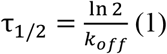

### Thermodynamic parameters of CYP143 and FdxE interaction

Sensorgrams were obtained at temperatures of 10, 15, 20, 25, 30, and 35°C. FdxE concentrations, the flow rate and contact time were the same as described above. The Gibbs free energy (ΔG) was calculated from the equation (2) using the Kd value obtained.

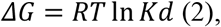

where T is the absolute temperature (°K), R - universal gas constant (J*mol^-1^*K^-1^), Kd - equilibrium dissociation constant of the protein-protein complex (M).

Enthalpy change (ΔH) was determined by plotting lnK_d_ versus 1000/T (Van ‘t Hoff plot) according to the linear form of the Van ‘t Hoff equation (3), using the slope of the linear regression line (ΔΗ/R).

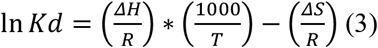

The entropy change (-TΔS) was calculated from the equation (4).

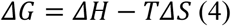

### Data collection, and X-ray structure determination

Diffraction data were collected at the European Synchrotron Radiation Facility (ESRF) beamlines ID23-1, ID30A1 and ID30B for CYP143, Fdx and FdxE−CYP143, respectively. The data collection strategy was optimized in BEST^43^,^44^. To increase data completeness for the CYP143 P1 crystals, two datasets from the same crystal were collected using different kappa angles (0° and 1°, respectively).

All data were processed in the XDS software package^45^. Processed data were corrected for anisotropy using the STARANISO server^46^ (http://staraniso.globalphasing.org/cgi-bin/staraniso.cgi).

A local mean I/σ(I) value of 0.50 was used to determine the anisotropic diffraction-limit surface. The phase problem for CYP143 was solved in the automatic molecular replacement pipeline MoRDa^47^, where structure of CYP101D2_Y96A_ (PDB ID: 4DXY^48^) was used as a starting model. The obtained space group was P1 with one molecule per asymmetric unit. Initially, molecular replacement gave model with poor-quality maps and the solution was further optimized in Morph model from Phenix^49,50^. The resultant model was than rebuilt using ARP/wARP web service^51,11^.

The phase problem for Fdx was solved in the automatic molecular replacement pipeline MoRDa^47^ from the CCP4 Online web service^52^, where structure of [4Fe-4S] ferredoxin (PDB ID: 1VJW^53^) was used as a starting model. The obtained space group was H32 with one molecule per asymmetric unit. Initially, molecular replacement gave model with poor-quality maps and the solution was further optimized in Morph model from Phenix^49,50^. The resultant model was then rebuilt in phenix.autobuild^51^.

To solve the phase problem for the FdxE−CYP143 complex, polyala models of the refined CYP143 and Fdx were used for molecular replacement in Phaser^54^. The space group for the complex was P1 and contained one molecule of both CYP143 and Fdx in ASU. The resultant model was then rebuilt in phenix.autobuild^55^.

For all the models multiple rounds of TLS-refinement^56^ were done in phenix.refine^57^ and refmac5^58^. Interactive refinement was performed in Coot^59^. The quality of the final models was analyzed using phenix.molprobity^60^. Fe ion in the Fdx structure was validated using Check My Metal web service^61^ and anomalous difference map, built with FFT from CCP4^62^. Data collection and final refinement statistics are presented in Table S1. Figures were rendered in PyMOL.

Structures of FdxE−CYP143, CYP143 and Fdx have been deposited in the Protein Data Bank (PDB) under the accession codes 8AMQ, 8AMO and 8AMP, respectively.

### Small-angle X-ray scattering measurements and data processing

SAXS measurements were carried on BM29 BioSAXS beamline (ESRF, Grenoble, France)^63^. All measurements were performed with 100% of beam intensity at a wavelength of 0.9918 Å (12.5 keV). Initial data processing was performed automatically using the EDNA pipeline^64,65^. SAXS profiles were obtained at a protein concentrations of 2.2 mg/ml, which was small enough to neglect the influence of the structure factor S(q) on the scattering curves^66,67^, in contract to systems with relatively high sample concentrations (∼1% or more ^68^). Exposure time was 8 s for the sample and 205 s for the buffer.

SAXS profile *I*(*q*) were processed using ATSAS^69^ software suite. The SAXS curve obtained for the fused complex FdxE−CYP143 has a wide Guinier region (Fig. S7A) that confirms that the protein has a globular structure with *R*_g_ = 25 Å. The behavior of the dimensionless Kratky plot (Fig. S7B) also confirms a similarity of the complex FdxE−CYP143 to a compact globular structure in solution. The plot has a maximum value of 1.120 at *qR*_g_ = 1.786, which is very close to the properties of the peak in the case of ideally globular particles (1.104 at *qR*_g_ = 3)^70^. A peak shift towards higher *qR*_g_ values could indicate partial disorder or an elongated protein shape^71,72^, but this was not observed in our data.

Distance distribution function *P*(*r*) (Fig. S7C) was calculated using the GNOM program. Experimental MW was calculated from Porod volume and as *V*_c_^2^/ 123.1 *R* ^73^. CORAL program^26^ was used to perform SAXS-based modeling of the residues of random loops missed in crystal structure of the fused complex FdxE−CYP143.

We used the EOM program^27^ from ATSAS online web platform to fit SAXS experimental data with an ensemble of proteins with variable positions of CYP143 and FdxE. Pool of the structures generated by EOM corresponds to the distribution of *R*_g_ from 23 Å to 45 Å. However, the experimental *R*_g_ = 25 Å (Table S1; Guinier plot is shown in Fig. S7A) is close to the left border of the pool distribution. The distribution of *R*_g_ for models selected by the genetic algorithm (Fig. S7B) has a maximum at ∼25 Å and is not symmetrical due to elongated right side wing. The other details of SAXS measurements and data treatment are presented in Table S1 (prepared in accordance with^74^) along with the data processing results. SAXS data was deposited with SASBDB (http://sasbdb.org)^75^ with accession code SASDPL2.

## Author contributions

A.G. helped conceive experiments, prepared figures, corrected the manuscript, discussed data. S.S. and A.K. performed molecular cloning. T.V. and A.K. purified and crystallized all proteins. T.S. and K.T. performed bioinformatic analysis. L.K., O.G., E.Y. and A.I. performed SPR measurements and analyzed data. E.M., S.B., K.K., A.M., V.B. collected crystallographic data. S.B., E.M., I.K., M.Sh. solved and refined crystallographic structures under supervision of V.B. Y.R., A.K. collected and treated SAXS data under supervision of V.G. and V.B. I.K. and I.O. performed binding assays. N.S. conceived and coordinated the project, wrote the manuscript, prepared figures, analyzed data and directed the bioinformatic analyses. All authors contributed to the writing of the Methods section.

## Data availability

Coordinates and structure factors for the structures of Fdx, CYP143 and the complex FdxE–CYP143 have been deposited in the Protein Data Bank (PDB) under the accession codes 8AMP, 8AMO and 8AMQ, respectively.

## Acknowledgements

The SPR analysis was done within the framework of the Program for Basic Research in the Russian Federation for a long-term period (2021–2030) (No 122030100168-2) using the equipment of “Human Proteome” Core Facility of the Institute of Biomedical Chemistry.

## Notes

### Competing Interest Statement

The authors have declared no competing interest.

## References

1. Beinert H, Holm RH, Munck E. Iron-sulfur clusters: nature’s modular, multipurpose structures. Science 277, 653–659 (1997).

2. Hall DO, Cammack R, Rao KK. Role for ferredoxins in the origin of life and biological evolution. Nature 233, 136–138 (1971).

3. Hall DO, Cammack R, Rao KK. The iron-sulphur proteins: Evolution of a ubiquitous protein from model systems to higher organisms. Origins of Life 5, 363–386 (1974).

4. Braymer JJ, Freibert SA, Rakwalska-Bange M, Lill R. Mechanistic concepts of iron-sulfur protein biogenesis in Biology. Biochim Biophys Acta Mol Cell Res 1868, 118863 (2021).

5. Hosseinzadeh P, Lu Y. Design and fine-tuning redox potentials of metalloproteins involved in electron transfer in bioenergetics. Biochim Biophys Acta 1857, 557–581 (2016).

6. Kim JY, Nakayama M, Toyota H, Kurisu G, Hase T. Structural and mutational studies of an electron transfer complex of maize sulfite reductase and ferredoxin. J Biochem 160, 101–109 (2016).

7. Watanabe T, Shima S. MvhB-type Polyferredoxin as an Electron-transfer Chain in Putative Redox-enzyme Complexes. Chemistry Letters 50, 353–360 (2021).

8. Meyer J. Iron-sulfur protein folds, iron-sulfur chemistry, and evolution. J Biol Inorg Chem 13, 157–170 (2008).

9. Nzuza N, Padayachee T, Chen W, Gront D, Nelson DR, Syed K. Diversification of Ferredoxins across Living Organisms. Curr Issues Mol Biol 43, 1374–1390 (2021).

10. Campbell IJ, Bennett GN, Silberg JJ. Evolutionary Relationships Between Low Potential Ferredoxin and Flavodoxin Electron Carriers. Front Energy Res 7, (2019).

11. Tilley GJ, Camba R, Burgess BK, Armstrong FA. Influence of electrochemical properties in determining the sensitivity of [4Fe-4S] clusters in proteins to oxidative damage. Biochemical Journal 360, 717–726 (2001).

12. Gao-Sheridan HS, et al. A T14C variant of Azotobacter vinelandii ferredoxin I undergoes facile [3Fe-4S]0 to [4Fe-4S]2+ conversion in vitro but not in vivo. J Biol Chem 273, 33692–33701 (1998).

13. Li S, Du L, Bernhardt R. Redox Partners: Function Modulators of Bacterial P450 Enzymes. Trends Microbiol 28, 445–454 (2020).

14. Nkosi BVZ, Padayachee T, Gront D, Nelson DR, Syed K. Contrasting Health Effects of Bacteroidetes and Firmicutes Lies in Their Genomes: Analysis of P450s, Ferredoxins, and Secondary Metabolite Clusters. Int J Mol Sci 23, (2022).

15. McLean KJ, et al. Biophysical characterization of the sterol demethylase P450 from Mycobacterium tuberculosis, its cognate ferredoxin, and their interactions. Biochemistry 45, 8427–8443 (2006).

16. Zanno A, Kwiatkowski N, Vaz AD, Guardiola-Diaz HM. MT FdR: a ferredoxin reductase from M. tuberculosis that couples to MT CYP51. Biochim Biophys Acta 1707, 157–169 (2005).

17. Lu Y, et al. Recombinant expression and biochemical characterization of Mycobacterium tuberculosis 3Fe-4S ferredoxin Rv1786. Appl Microbiol Biotechnol 101, 7201–7212 (2017).

18. Podust LM, Poulos TL, Waterman MR. Crystal structure of cytochrome P450 14alpha -sterol demethylase (CYP51) from Mycobacterium tuberculosis in complex with azole inhibitors. Proc Natl Acad Sci U S A 98, 3068–3073 (2001).

19. Kissinger CR, Sieker LC, Adman ET, Jensen LH. Refined crystal structure of ferredoxin II from Desulfovibrio gigas at 1·7 Å. Journal of Molecular Biology 219, 693–715 (1991).

20. Dehal PS, et al. MicrobesOnline: an integrated portal for comparative and functional genomics. Nucleic Acids Res 38, D396–400 (2010).

21. Groschel MI, Sayes F, Simeone R, Majlessi L, Brosch R. ESX secretion systems: mycobacterial evolution to counter host immunity. Nat Rev Microbiol 14, 677–691 (2016).

22. Schreiber G, Haran G, Zhou HX. Fundamental aspects of protein-protein association kinetics. Chem Rev 109, 839–860 (2009).

23. Stites WE. Proteinminus signProtein Interactions: Interface Structure, Binding Thermodynamics, and Mutational Analysis. Chem Rev 97, 1233–1250 (1997).

24. Brady GP, Sharp KA. Entropy in protein folding and in protein—protein interactions. Current Opinion in Structural Biology 7, 215–221 (1997).

25. Strushkevich N, MacKenzie F, Cherkesova T, Grabovec I, Usanov S, Park HW. Structural basis for pregnenolone biosynthesis by the mitochondrial monooxygenase system. Proc Natl Acad Sci U S A 108, 10139–10143 (2011).

26. Petoukhov MV, et al. New developments in the ATSAS program package for small-angle scattering data analysis. J Appl Crystallogr 45, 342–350 (2012).

27. Tria G, Mertens HD, Kachala M, Svergun DI. Advanced ensemble modelling of flexible macromolecules using X-ray solution scattering. IUCrJ 2, 207–217 (2015).

28. Khramtsov YV, et al. Low-resolution structures of modular nanotransporters shed light on their functional activity. Acta Crystallogr D Struct Biol 76, 1270–1279 (2020).

29. Child SA, Bradley JM, Pukala TL, Svistunenko DA, Le Brun NE, Bell SG. Electron transfer ferredoxins with unusual cluster binding motifs support secondary metabolism in many bacteria. Chem Sci 9, 7948–7957 (2018).

30. O’Keefe DP, et al. Ferredoxins from two sulfonylurea herbicide monooxygenase systems in Streptomyces griseolus. Biochemistry 30, 447–455 (1991).

31. Bell SG, Hoskins N, Xu F, Caprotti D, Rao Z, Wong LL. Cytochrome P450 enzymes from the metabolically diverse bacterium Rhodopseudomonas palustris. Biochem Biophys Res Commun 342, 191–196 (2006).

32. Zhang T, Zhang A, Bell SG, Wong LL, Zhou W. The structure of a novel electron-transfer ferredoxin from Rhodopseudomonas palustris HaA2 which contains a histidine residue in its iron-sulfur cluster-binding motif. Acta Crystallogr D Biol Crystallogr 70, 1453–1464 (2014).

33. Lamb DC, et al. Concerning P450 Evolution: Structural Analyses Support Bacterial Origin of Sterol 14alpha-Demethylases. Mol Biol Evol 38, 952–967 (2021).

34. Jackson CJ, et al. A novel sterol 14alpha-demethylase/ferredoxin fusion protein (MCCYP51FX) from Methylococcus capsulatus represents a new class of the cytochrome P450 superfamily. J Biol Chem 277, 46959–46965 (2002).

35. Hargrove TY, Lamb DC, Smith JA, Wawrzak Z, Kelly SL, Lepesheva GI. Unravelling the role of transient redox partner complexes in P450 electron transfer mechanics. Scientific Reports 12, (2022).

36. Lin H, et al. Mercury methylation by metabolically versatile and cosmopolitan marine bacteria. ISME J 15, 1810–1825 (2021).

37. Zhang L, et al. Structural insight into the electron transfer pathway of a self-sufficient P450 monooxygenase. Nat Commun 11, 2676 (2020).

38. Wang Z, Shaik S, Wang B. Conformational Motion of Ferredoxin Enables Efficient Electron Transfer to Heme in the Full-Length P450TT. J Am Chem Soc 143, 1005–1016 (2021).

39. Neeli R, et al. The dimeric form of flavocytochrome P450 BM3 is catalytically functional as a fatty acid hydroxylase. FEBS Lett 579, 5582–5588 (2005).

40. Mirdita M, Schutze K, Moriwaki Y, Heo L, Ovchinnikov S, Steinegger M. ColabFold: making protein folding accessible to all. Nat Methods 19, 679–682 (2022).

41. Mohamed H, Child SA, Bruning JB, Bell SG. A comparison of the bacterial CYP51 cytochrome P450 enzymes from Mycobacterium marinum and Mycobacterium tuberculosis. J Steroid Biochem Mol Biol 221, 106097 (2022).

42. Ortega Ugalde S, et al. Linking cytochrome P450 enzymes from Mycobacterium tuberculosis to their cognate ferredoxin partners. Appl Microbiol Biotechnol 102, 9231–9242 (2018).

43. Popov AN, Bourenkov GP. Choice of data-collection parameters based on statistic modelling. Acta Crystallogr D Biol Crystallogr 59, 1145–1153 (2003).

44. Bourenkov GP, Popov AN. A quantitative approach to data-collection strategies. Acta Crystallogr D Biol Crystallogr 62, 58–64 (2006).

45. Kabsch W. Xds. Acta Crystallogr D Biol Crystallogr 66, 125–132 (2010).

46. Vonrhein C, et al. Advances in automated data analysis and processing within autoPROC, combined with improved characterisation, mitigation and visualisation of the anisotropy of diffraction limits using STARANISO. Acta Crystallographica Section A 74, a360 (2018).

47. Vagin A, Lebedev A. MoRDa, an automatic molecular replacement pipeline. Acta Crystallographica Section A 71, s19 (2015).

48. Bell SG, Yang W, Dale A, Zhou W, Wong LL. Improving the affinity and activity of CYP101D2 for hydrophobic substrates. Appl Microbiol Biotechnol 97, 3979–3990 (2013).

49. Terwilliger TC, et al. Improved crystallographic models through iterated local density-guided model deformation and reciprocal-space refinement. Acta Crystallogr D Biol Crystallogr 68, 861–870 (2012).

50. Liebschner D, et al. Macromolecular structure determination using X-rays, neutrons and electrons: recent developments in Phenix. Acta Crystallogr D Struct Biol 75, 861–877 (2019).

51. Chojnowski G, et al. The use of local structural similarity of distant homologues for crystallographic model building from a molecular-replacement solution. Acta Crystallogr D Struct Biol 76, 248–260 (2020).

52. Krissinel E, Uski V, Lebedev A, Winn M, Ballard C. Distributed computing for macromolecular crystallography. Acta Crystallogr D Struct Biol 74, 143–151 (2018).

53. Macedo-Ribeiro S, Darimont B, Sterner R, Huber R. Small structural changes account for the high thermostability of 1[4Fe–4S] ferredoxin from the hyperthermophilic bacterium Thermotoga maritima. Structure 4, 1291–1301 (1996).

54. McCoy AJ, Grosse-Kunstleve RW, Adams PD, Winn MD, Storoni LC, Read RJ. Phaser crystallographic software. J Appl Crystallogr 40, 658–674 (2007).

55. Terwilliger TC, et al. Iterative model building, structure refinement and density modification with the PHENIX AutoBuild wizard. Acta Crystallogr D Biol Crystallogr 64, 61–69 (2008).

56. Painter J, Merritt EA. Optimal description of a protein structure in terms of multiple groups undergoing TLS motion. Acta Crystallogr D Biol Crystallogr 62, 439–450 (2006).

57. Afonine PV, et al. Towards automated crystallographic structure refinement with phenix.refine. Acta Crystallogr D Biol Crystallogr 68, 352–367 (2012).

58. Murshudov GN, et al. REFMAC5 for the refinement of macromolecular crystal structures. Acta Crystallogr D Biol Crystallogr 67, 355–367 (2011).

59. Emsley P, Lohkamp B, Scott WG, Cowtan K. Features and development of Coot. Acta Crystallogr D Biol Crystallogr 66, 486–501 (2010).

60. Williams CJ, et al. MolProbity: More and better reference data for improved all-atom structure validation. Protein Sci 27, 293–315 (2018).

61. Handing KB, Niedzialkowska E, Shabalin IG, Kuhn ML, Zheng H, Minor W. Characterizing metal-binding sites in proteins with X-ray crystallography. Nat Protoc 13, 1062–1090 (2018).

62. Winn MD, et al. Overview of the CCP4 suite and current developments. Acta Crystallogr D Biol Crystallogr 67, 235–242 (2011).

63. Pernot P, et al. Upgraded ESRF BM29 beamline for SAXS on macromolecules in solution. J Synchrotron Radiat 20, 660–664 (2013).

64. Brennich ME, et al. Online data analysis at the ESRF bioSAXS beamline, BM29. Journal of Applied Crystallography 49, 203–212 (2016).

65. Incardona MF, Bourenkov GP, Levik K, Pieritz RA, Popov AN, Svensson O. EDNA: a framework for plugin-based applications applied to X-ray experiment online data analysis. J Synchrotron Radiat 16, 872–879 (2009).

66. Poster Sessions. FEBS Journal 281, 65–784 (2014).

67. Zabelskii DV, et al. Ambiguities and completeness of SAS data analysis: investigations of apoferritin by SAXS/SANS EID and SEC-SAXS methods. Journal of Physics: Conference Series 994, 012017 (2018).

68. Murugova TN, et al. Mechanisms of membrane protein crystallization in ‘bicelles’. Sci Rep 12, 11109 (2022).

69. Franke D, et al. ATSAS 2.8: a comprehensive data analysis suite for small-angle scattering from macromolecular solutions. J Appl Crystallogr 50, 1212–1225 (2017).

70. Burger VM, Arenas DJ, Stultz CM. A Structure-free Method for Quantifying Conformational Flexibility in proteins. Sci Rep 6, 29040 (2016).

71. Ryzhykau YL, et al. Molecular model of a sensor of two-component signaling system. Sci Rep 11, 10774 (2021).

72. Ryzhykau YL, et al. Ambiguities in and completeness of SAS data analysis of membrane proteins: the case of the sensory rhodopsin II-transducer complex. Acta Crystallogr D Struct Biol 77, 1386–1400 (2021).

73. Rambo RP, Tainer JA. Accurate assessment of mass, models and resolution by small-angle scattering. Nature 496, 477–481 (2013).

74. Trewhella J, et al. 2017 publication guidelines for structural modelling of small-angle scattering data from biomolecules in solution: an update. Acta Crystallogr D Struct Biol 73, 710–728 (2017).

75. Kikhney AG, Borges CR, Molodenskiy DS, Jeffries CM, Svergun DI. SASBDB: Towards an automatically curated and validated repository for biological scattering data. Protein Sci 29, 66–75 (2020).

