## Supplemental Information for "Structural insights into 3Fe-4S ferredoxins diversity in *M.tuberculosis* highlighted by a first redox complex with P450"

**Supplementary materials:**
**Fig. S1**. The geometry of [3Fe-4S] cluster (the cluster plus the 3 cysteine Sγ ligands) (A) and values of individual bond lengths and angles for Fdx and FdxE and other structurally studied ferredoxins (B).

A

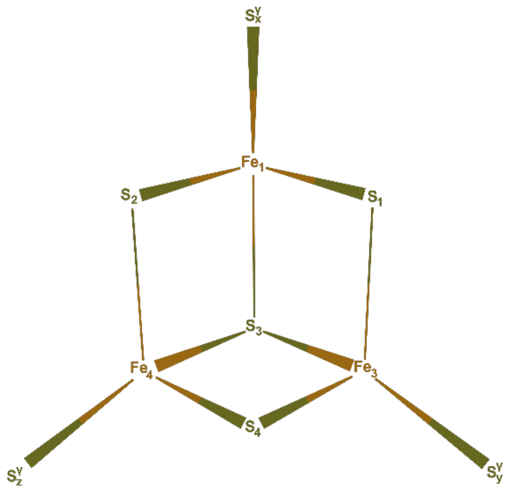


B

| PDB ID | $Fe_1$-$S_x^{\gamma}$ | $Fe_3$-$S_y^{\gamma}$ | $Fe_4$-$S_z^{\gamma}$ | $Fe_1$-$S_1$ | $Fe_1$-$S_2$ | $Fe_1$-$S_3$ | $Fe_3$-$S_1$ | $Fe_3$-$S_3$ | $Fe_3$-$S_4$ | $Fe_4$-$S_2$ | $Fe_4$-$S_3$ | $Fe_4$-$S_4$ |
|---|---|---|---|---|---|---|---|---|---|---|---|---|
| Fdx | 2.32 | 2.32 | 2.31 | 2.25 | 2.25 | 2.25 | 2.26 | 2.25 | 2.25 | 2.26 | 2.26 | 2.25 |
| FdxE | 2.32 | 2.35 | 2.34 | 2.27 | 2.30 | 2.25 | 2.27 | 2.29 | 2.29 | 2.27 | 2.29 | 2.29 |
| 1FXD | 2.29 | 2.28 | 2.22 | 2.23 | 2.33 | 2.32 | 2.26 | 2.32 | 2.27 | 2.29 | 2.31 | 2.22 |
| $1SJ1_a$ | 2.28 | 2.32 | 2.23 | 2.17 | 2.32 | 2.33 | 2.25 | 2.31 | 2.26 | 2.34 | 2.38 | 2.24 |
| $1SJ1_b$ | 2.27 | 2.32 | 2.28 | 2.23 | 2.32 | 2.35 | 2.33 | 2.36 | 2.23 | 2.32 | 2.31 | 2.32 |
| 4ID8 | 2.31 | 2.25 | 2.19 | 2.21 | 2.22 | 2.19 | 2.21 | 2.14 | 2.24 | 2.27 | 2.16 | 2.21 |
| 4OV1 | 2.39 | 2.30 | 2.25 | 2.21 | 2.19 | 2.23 | 2.22 | 2.20 | 2.20 | 2.23 | 2.21 | 2.21 |

| PDB ID | $S_x^{\gamma}$-$Fe_1$-$S_1$ | $S_x^{\gamma}$-$Fe_1$-$S_2$ | $S_x^{\gamma}$-$Fe_1$-$S_3$ | $S_y^{\gamma}$-$Fe_3$-$S_1$ | $S_y^{\gamma}$-$Fe_3$-$S_3$ | $S_y^{\gamma}$-$Fe_3$-$S_4$ | $S_z^{\gamma}$-$Fe_4$-$S_2$ | $S_z^{\gamma}$-$Fe_4$-$S_3$ | $S_z^{\gamma}$-$Fe_4$-$S_4$ |
|---|---|---|---|---|---|---|---|---|---|
| Fdx | 101.8 | 114.8 | 116.5 | 104.9 | 120.1 | 110.8 | 117.5 | 102.8 | 116.9 |
| FdxE | 104.9 | 109.7 | 120.4 | 110.3 | 119.3 | 105.5 | 108.4 | 109.1 | 111.7 |
| 1FXD | 106.0 | 114.0 | 118.8 | 110.5 | 116.4 | 116.0 | 110.4 | 110.5 | 121.4 |
| $1SJ1_a$ | 104.7 | 112.9 | 115.3 | 109.9 | 116.6 | 111.3 | 109.8 | 112.1 | 117.6 |
| $1SJ1_b$ | 104.2 | 114.0 | 115.9 | 107.9 | 116.1 | 113.0 | 111.0 | 111.6 | 115.6 |
| 4ID8 | 104.4 | 114.6 | 115.6 | 104.0 | 119.5 | 112.9 | 102.8 | 116.8 | 116.8 |
| 4OV1 | 111.7 | 112.4 | 110.3 | 103.7 | 122.5 | 111.7 | 105.3 | 112.4 | 117.7 |

| PDB ID | $S_1$-$Fe_1$-$S_2$ | $S_1$-$Fe_1$-$S_3$ | $S_2$-$Fe_1$-$S_3$ | $S_1$-$Fe_3$-$S_3$ | $S_1$-$Fe_3$-$S_4$ | $S_3$-$Fe_3$-$S_4$ | $S_2$-$Fe_4$-$S_3$ | $S_2$-$Fe_4$-$S_4$ | $S_3$-$Fe_4$-$S_4$ |
|---|---|---|---|---|---|---|---|---|---|
| Fdx | 111.1 | 107.0 | 105.5 | 106.8 | 108.4 | 105.4 | 105.2 | 107.5 | 105.4 |
| FdxE | 113.7 | 103.1 | 105.2 | 101.8 | 114.3 | 106.0 | 104.8 | 116.5 | 105.9 |
| 1FXD | 109.2 | 105.1 | 103.3 | 104.2 | 107.2 | 101.3 | 104.8 | 105.0 | 103.4 |
| $1SJ1_a$ | 111.2 | 105.3 | 107.3 | 103.4 | 112.0 | 103.3 | 105.0 | 109.8 | 101.5 |
| $1SJ1_b$ | 109.6 | 106.7 | 106.1 | 103.5 | 111.7 | 104.4 | 107.4 | 107.6 | 103.1 |

| | | | | | | | | | |
|---|---|---|---|---|---|---|---|---|---|
| 4ID8 | 112.2 | 103.8 | 105.8 | 105.4 | 108.5 | 105.8 | 104.8 | 108.4 | 106.3 |
| 4OV1 | 119.3 | 101.2 | 100.3 | 101.6 | 114.1 | 103.2 | 99.9 | 117.6 | 102.7 |

| PDB ID | $Fe_1$-$S_1$- $Fe_3$ | $Fe_1$-$S_3$- $Fe_3$ | $Fe_3$-$S_3$- $Fe_4$ | $Fe_3$-$S_4$- $Fe_4$ | $Fe_4$-$S_2$- $Fe_1$ | $Fe_4$-$S_3$- $Fe_1$ |
|---|---|---|---|---|---|---|
| Fdx | 69.5 | 69.6 | 71.6 | 71.7 | 70.9 | 71.0 |
| FdxE | 73.1 | 73.2 | 69.9 | 69.9 | 70.3 | 70.8 |
| 1FXD | 74.0 | 71.0 | 73.9 | 76.7 | 73.4 | 72.9 |
| $1SJ1_a$ | 73.6 | 69.7 | 71.9 | 75.4 | 71.1 | 70.3 |
| $1SJ1_b$ | 72.3 | 69.8 | 71.3 | 73.5 | 70.6 | 70.2 |
| 4ID8 | 70.8 | 72.7 | 72.7 | 70.0 | 69.9 | 72.5 |
| 4OV1 | 74.7 | 74.6 | 73.8 | 73.9 | 74.8 | 74.5 |

**Fig. S2.** Species name and protein ID (in parenthesis) for ferredoxin sequences in Fig.2.

**mav_1 -** M.avium (WP_062890415.1); **mma_1 -** M.marinum (WP_117428654.1; **msm_1 -** M.smegmatis (WP_003896201.1); **msp_1** – Mycobacterium sp. (WP_009953936.1; **afe_1** –Acidimicrobium ferrooxidans (WP_015798421.1); **aro_1** – Actinospica robiniae (WP_034260932.1); **aau_1** –Arthrobacter aurescens (WP_011777263.1); **cac_1** – Catenulispora acidiphila (WP_015795994.1); **fsp_1** - Frankia_sp. (WP_020464245.1); **faf_1** – Mycobacterium sp. (WP_011724409.1); **nha_1-** Nitrobacter hamburgensis (WP_011511137.1); **nsp_1a -** Nitrobacter_sp._Nb311A (WP_009800054.1); **nsp_1b** - Nocardioides_sp. (WP_011757263.1); **nar_1** – Novosphingobium aromaticivorans (WP_011906924.1); **pae_1** – Pseudomonas aeruginosa (WP_121351230.1); **reu_1** - Ralstonia eutropha (WP_136227872.1); **ret_1** – Rhizobium etli (AAM54837.2); **rop_1** – Rhodococcus opacus (WP_015890824.1); **rpa_1** – Rhodopseudomonas palustris (WP_011157460.1); **swi_1 -** Sphingomonas wittichii (WP_011952583.1); **sco_1** – Streptomyces coelicolor (AGO88622.1); **tfu_1** – Thermobifida fusca (WP_011291910.1); **tcu_1 -** Thermomonospora curvata (WP_012851129.1); **mab_2a** - M.abscessus (WP_005059330.1); **mav_2a -** M.avium (WP_003875758.1); **mma_2 -** M.ulcerans Agy99 (ABL03175.1); **mtb_2 -** M.tuberculosis T85 (EFD76427.1); **mva_2 -** Mycolicibacterium sp. (WP_011782303.1); **mab_2b -** Mycobacteroides sp. (WP_180748986.1); **mav_2b -** M.avium (WP_224188525.1); **mav_2c -** M.avium (WP_003873007.1); **msm_2 -** M.smegmatis (WP_011730111.1); **msp_2** – Mycolicibacterium monacense (WP_011768425.1); **mul_2 -** M.marinum (WP_094360996.1); **afe_2** – Acidimicrobium ferrooxidans (WP_015798014.1); **fsp_2a** – Frankia sp. (WP_011435700.1); **fsp_2b -** Frankia_sp. (WP_071060280.1); **gbr_2** - Gordonia_bronchialis (WP_012835523.1); **hoc_2** –Haliangium ochraceum (WP_012827286.1); **kfl_2** – Kribbella flavida (WP_012921518.1); **kra_2** – Ktedonobacter racemifer (WP_007922270.1); **mca_2** - Methylococcus capsulatus (WP_010961920.1); **mxa_2 -** Myxococcus xanthus (DK 1622_ABF90123.1); **nfa_2** – Nocardia sp. (WP_011207065.1); **nsp_2** – Nocardioides sp. (WP_011756255.1); **rer_2** – Rhodococcus erythropolis (WP_073511998.1); **rop_2** – Rhodococcus opacus (WP_015889158.1); **str_2 –** Salinispora tropica (WP_012014024.1); **sav_2** – Streptomyces sp. (WP_010982020.1); **sro_2** – Streptosporangium sp. (WP_012895195.1)

**Fig. S3.** Gene context for Rv076c (A) and Rv1786 (B). Genes colored according to their homology group; non-related genes are in grey. COGs - cluster of orthologous groups. The gene context was generated using MicrobesOnline [doi: 10.1093/nar/gkp919].

A

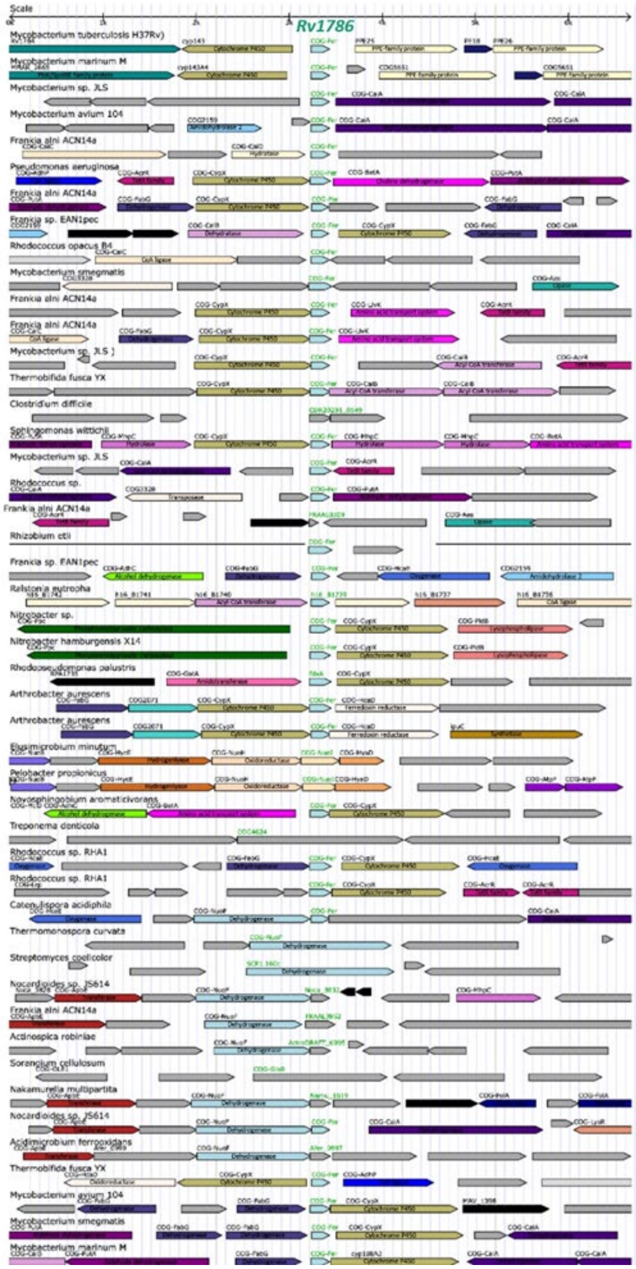


B

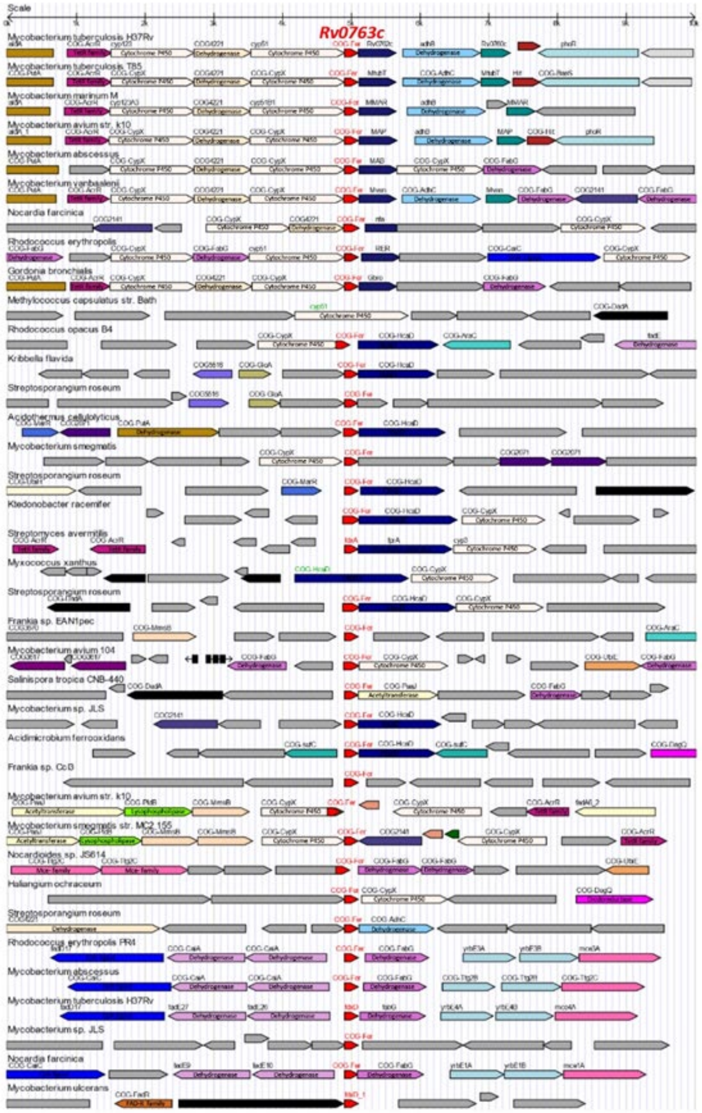

**Fig. S4**. Typical sensorgrams of interactions between immobilized on SA chip biotinylated CYP143 and FdxE in PBS buffer at 25 °C. Rv1786 in the following concentrations: 1 – 5 nM, 2 – 10 nM, 3 – 25 nM, 4 – 50 nM, 5 – 75 nM.

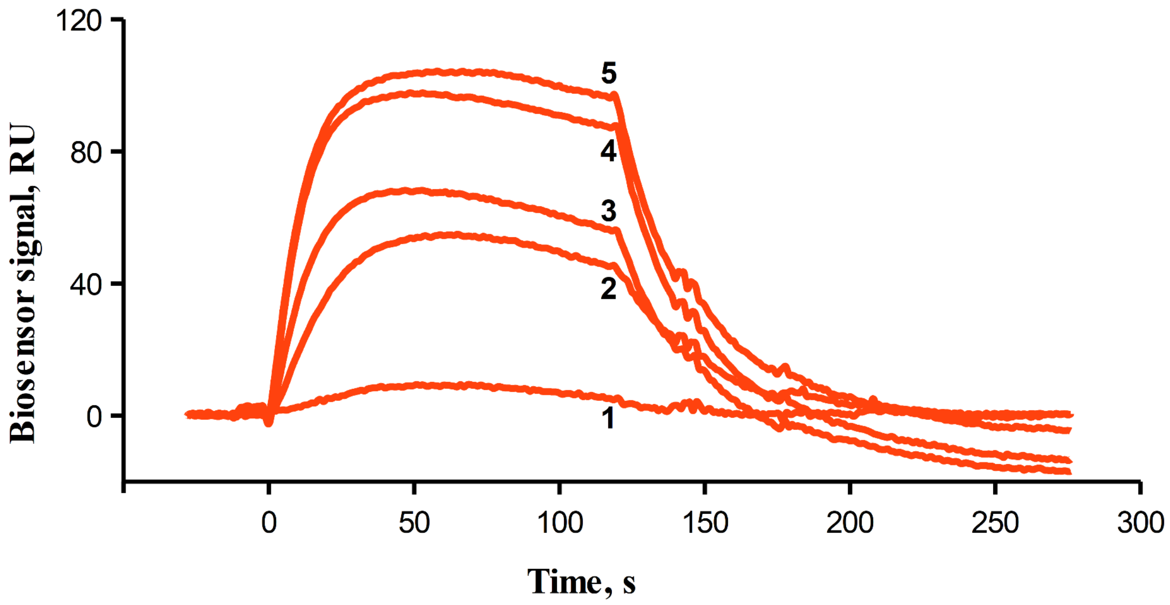

**Fig. S5.** The electrostatic surface potentials of the interaction faces of CYP143 (A) and FdxE (B). Negatively and positively charged surface areas are coloured red and blue, respectively. Residues that are involved in the interactions are labeled.

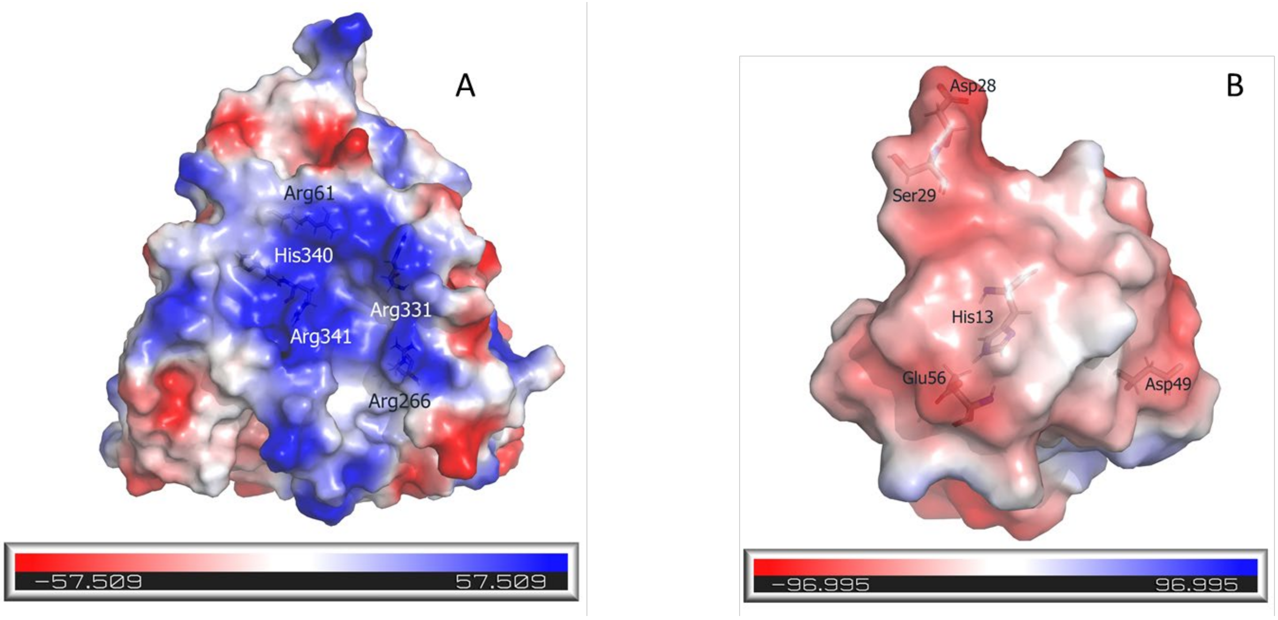

**Fig. S6. Results of SAXS data analysis for the FdxE–CYP143 complex.** A – SAXS $I(q)$ profile (orange circles) and their approximations corresponding to the PDB models obtained by using CORAL (purple dashed line) and EOM (blue solid line); B –$R_g$ distributions for the initial pool (gray histogram) and for a set of selected conformations (blue histogram).

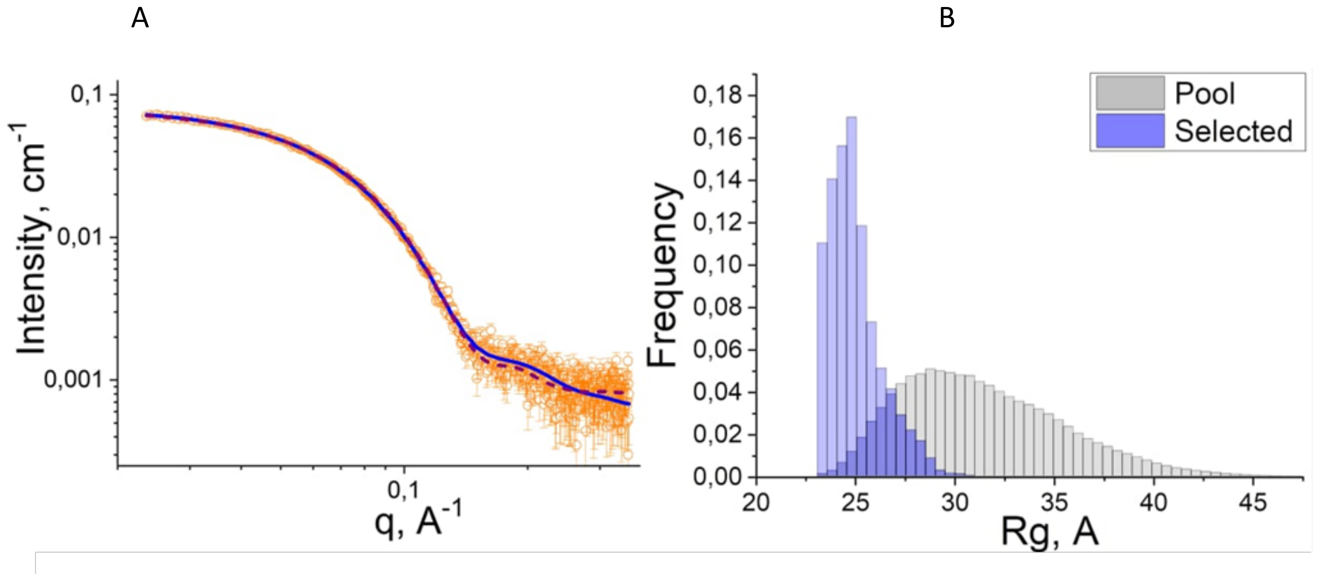


**Fig. S7. Analysis of SAXS data for the FdxE–CYP143 complex.** ***(a)*** – Guinier approximation. ***(b)*** – normalized Kratky plot (dashed lines are drawn at $qR_g = \sqrt{3}$ and $(qR_g)^2\, I(q) / I(0) = 1.104$). ***(c)*** – pair distance distribution function $P(r)$.

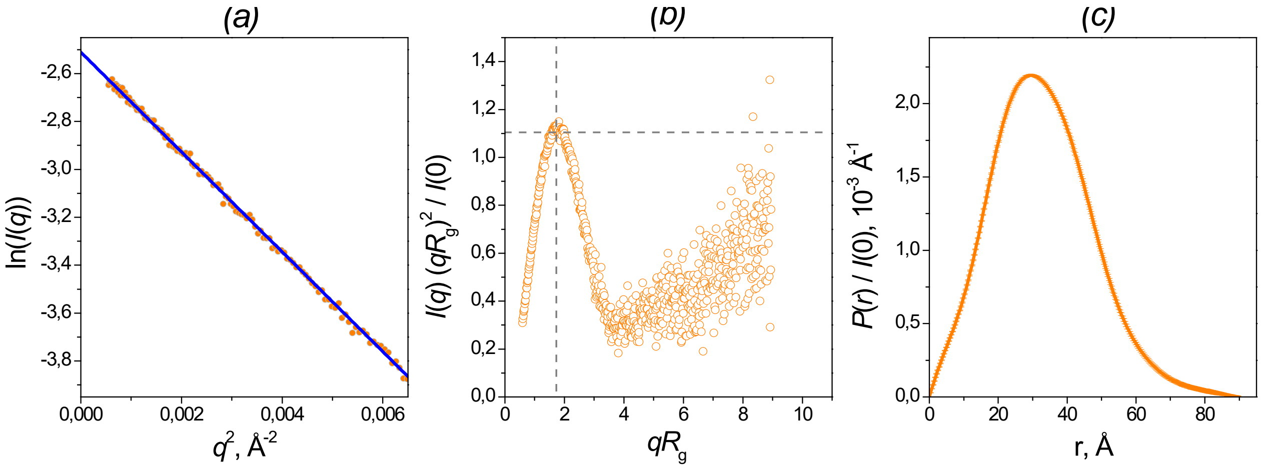

**Fig. S8. Summary of the Rv0763c and Rv1786 gene expression.** Constructed using http://www.microbesonline.org/

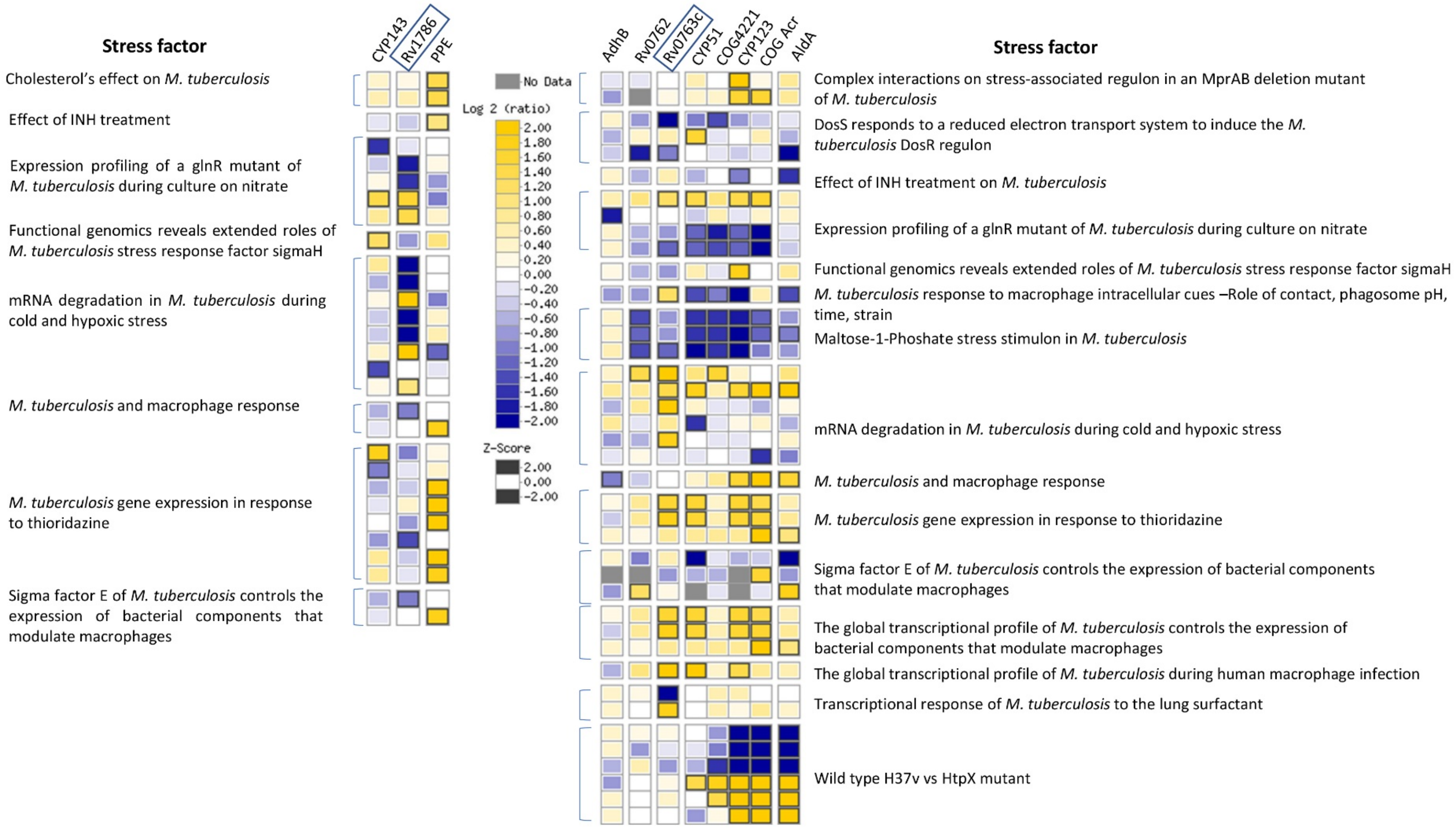

**Table S1.** MX data collection and refinement statistics

| Structure | CYP143 | | Fdx | | FdxE–CYP143 | |
|---|---|---|---|---|---|---|
| PDB ID | 8AMO | | 8AMP | | 8AMQ | |
| **Data collection** | | | | | | |
| | XSCALE | Staraniso | XSCALE | Staraniso | XSCALE | Staraniso |
| Beamline | ESRF ID23-1 | | ESRF ID30A1 | | ESRF ID30B | |
| Wavelength, Å | 0.972 | | 0.966 | | 0.976 | |
| Space group | P 1 | | H 32 | | P 1 | |
| Unit cell | 42.21, 48.53, 54.15, 111.65, 99.50, 109.34 | | 46.96, 46.96, 162.67, 90, 90, 120 | | 53.20, 54.35, 69.04, 67.71, 77.21, 61.63 | |
| Resolution range (Å)* | 27.78 - 1.40 (1.45 - 1.40) | | 28.76 - 2.00 (2.05 - 2.00) | | 45.32 - 1.60 (1.67 - 1.60) | |
| Resolution limits (Å) | 1.4 | 1.36, 1.60, 1.39 | 2.0 | 1.65, 1.65, 2.57 | 1.6 | 1.55, 1.57, 1.90 |
| No. of total reflections | 469759 (33483) | 401442 (19716) | 46885 (3417) | 46875 (2491) | 274784 (17758) | 232292 (11410) |
| No. of unique reflections | 62494 (4500) | 53501 (2475) | 4958 (346) | 5093 (392) | 78791 (5469) | 66216 (3311) |
| Multiplicity | 7.5 (7.4) | 7.5 (7.4) | 9.5 (9.9) | 9.2 (6.4) | 3.5 (3.4) | 3.5 (3.4) |
| Completeness | | | | | | |
| spherical (%) | 88.8 (86.6) | 76.3 (35.5) | 99.9 (100.0) | 60.6 (17.2) | 94.8 (89.0) | 79.8 (32.7) |
| ellipsoidal (%) | - | 84.6 (71.5) | - | 91.5 (58.3) | - | 91.2 (72.5) |
| Mean I/sigma(I) | 14.0 (0.2) | 15.7 (0.8) | 11.0 (0.9) | 11.0 (0.4) | 4.7 (0.3) | 5.2 (0.8) |
| R-pim | 0.025 (3.771) | 0.023 (1.167) | 0.032 (0.880) | 0.034 (1.768) | 0.078 (1.638) | 0.081 (0.997) |
| CC1/2 | 1.000 (0.243) | 0.999 (0.329) | 0.998 (0.526) | 0.998 (0.127) | 0.994 (0.12) | 0.992 (0.249) |
| **Refinement** | | | | | | |
| Resolution range (Å) | 27.78 – 1.40 | | 28.76 – 2.00 | | 45.32 – 1.60 | |
| Reflections used in refinement | 53378 | | 4119 | | 66205 | |
| Reflections used for R-free | 2654 (5%) | | 416 (10%) | | 3304 (5%) | |
| R-work/R-free | 0.1684/0.1980 | | 0.2465/0.2859 | | 0.1773/0.2057 | |
| No. of non-hydrogen atoms | 3895 | | 487 | | 4479 | |
| macromolecules | 3304 | | 474 | | 3928 | |
| heme | 43 | | - | | 43 | |
| [3Fe-4S] | - | | 7 | | 7 | |
| solvent | 548 | | 6 | | 501 | |
| No. of protein residues | 385 | | 65 | | 464 | |
| RMS | | | | | | |
| bonds (Å) | 0.006 | | 0.005 | | 0.011 | |
| angles (°) | 0.90 | | 1.02 | | 1.37 | |
| Ramachandran favored (%) | 98.69 | | 100.00 | | 98.25 | |
| Ramachandran outliers (%) | 0.00 | | 0.00 | | 0.00 | |
| Average B-factor | 23.86 | | 56.80 | | 29.95 | |
| macromolecules | 22.65 | | 57.15 | | 29.30 | |
| heme | 15.36 | | - | | 15.82 | |
| [3Fe-4S] | - | | 54.86 | | 20.90 | |
| solvent | 31.83 | | 31.20 | | 36.39 | |
| No. of TLS groups | 3 | | 1 | | 4 | |

*Statistics for the highest-resolution shell are shown in parentheses

**Table S2.** SAXS experimental details and data evaluation summary.

| | |
|---|---|
| ***(a)* Sample details** | |
| Description of sequence | CYP143-Rv1786<br>His-tagged fused complex of cytochrome P450 143 (UniProt ID: P9WPL3) and ferredoxin Rv1786 (UniProt ID: O53937) from *Mycobacterium tuberculosis* |
| Extinction coefficient ε (A280, 0.1% $cm^{-1}$) [1] | 0.890 |
| Partial specific volume ν ($cm^3$ $g^{-1}$) [1] | 0.729 |
| Mean solute and solvent SLD ($10^{-6}$ $Å^{-2}$) [1] | 12.50, 9.75 |
| Mean scattering contrast Δρ ($10^{-6}$ $Å^{-2}$) [1] | 2.75 |
| Molecular mass (kDa) [1] | 53.561 |
| Sample concentration (mg $ml^{-1}$) | 2.2 |
| Solvent composition | 300 mM NaCl, 50 mM Tris/TrisHCl (pH 7.4), 10% glycerol |
| **(*b*) SAS data collection parameters** | |
| Instrument | ESRF BM29 |
| Wavelength (Å) | 0.9918 |
| Beam geometry (size, sample-to-detector distance) | 700 × 700 $\mu m^2$, 2.864 m |
| Sample configuration | 1.8 mm-diameter quartz capillary |
| *q*-measurement range ($Å^{-1}$) | 0.004 – 0.495 |
| Absolute scaling method | Comparison with scattering from pure $H_2O$ |
| Basis for normalization to constant counts | To transmitted intensity by direct beam counter |
| Exposure time, number of exposures | 2 frames/sec, 16 frames |
| Sample configuration including path length and flow rate | Sample was exposed to X-rays while flowing through the 1.8 mm-diameter quartz capillary. |
| Sample temperature (°C) | 20 |
| **(*c*) Software employed for SAS data reduction, analysis and interpretation** | |
| SAS data averaging and subtraction | PRIMUS from ATSAS 2.8.4 |
| Calculation of ε from sequence and $\underline{\nu}$ values from chemical composition | Peptide Property Calculator: http://biotools.nubic.northwestern.edu/proteincalc.html |
| Calculation of $\underline{\rho}$ values from chemical composition | SLD calculator: http://www.ncnr.nist.gov/resources/activation/ |
| Guinier, *P*(*r*) | GNOM 5.0 from ATSAS 3.0.5 |
| Atomic structure modelling | CORAL and EOM from ATSAS 3.0.1 (ATSAS online) |
| Molecular graphics | UCSF Chimera 1.16 |
| **(*d*) Structural parameters** | |
| Guinier analysis | |
| *I*(0) ($cm^{-1}$) | 0.0812 ± 0.0002 |
| $R_g$ (Å) | 25.0 ± 0.12 |
| *q*-range ($Å^{-1}$) ($qR_g$ range) | 0.0236 – 0.0518 (0.59 – 1.29) |
| *P*(*r*) analysis | |
| *I*(0) ($cm^{-1}$) | 0.0818 ± 0.0002 |
| $R_g$ (Å) | 25.38 ± 0.11 |
| $d_{max}$ (Å) | 90 |
| *q*-range ($Å^{-1}$) | 0.0236 – 0.3528 |
| $q_{min}$ $d_{max}$ / π | 0.676 |
| Total quality estimate (GNOM) | 0.915 |
| Volume ($V_P$) ($Å^3$) | 85035 |
| Experimental MW from $V_P$, kDa | 62.0 |
| Experimental MW = $V_c^2$ / 123.1 $R_g$, kDa [2] | 53 (± 10%) |

| | |
|---|---|
| ***(e)* Atomistic modelling** | |
| Method | SAXS-based rigid body modeling of complexes (CORAL) |
| $q$-range for fitting | 0.0236 – 0.3528 |
| Symmetry assumptions | P1 |
| Background subtraction ($cm^{-1}$) | 0.0007848 |
| $\chi^2$ value | 1.411 |
| Method | Ensemble Optimization Method (EOM) |
| $q$-range for fitting | 0.0236 – 0.3528 |
| Symmetry assumptions | P1 |
| Background subtraction ($cm^{-1}$) | 0.000 |
| $\chi^2$ value | 1.157 |
| $R_g$ values (Å), $d_{max}$ values (Å), and weights for multi-state model | 24.14, 84.90, ~0.20 (1/5)<br>27.32, 97.72, ~0.20 (1/5)<br>24.98, 89.97, ~0.20 (1/5)<br>24.34, 80.67, ~0.20 (1/5)<br>24.92, 83.70, ~0.20 (1/5) |
| Final ensemble $R_g$ (Å) and $d_{max}$ (Å) | 25.14, 87.39 |
| ***(f)* Data and model deposition IDs** | |
| | CYP143-rv1786<br>SASDPL2 |

[1] These values are calculated without taking into account ligands: heme for CYP143 and 3Fe-4S for FdxE.

[2] MW is calculated as $V_c^2$ / 123.1 $R_g$ according to [10.1038/nature12070].
